# Transgenic mice expressing human alpha-synuclein in noradrenergic neurons develop locus coeruleus pathology and non-motor features of Parkinson’s disease

**DOI:** 10.1101/857987

**Authors:** LM Butkovich, MC Houser, T Chalermpalanupap, KA Porter-Stransky, AF Iannitelli, JS Boles, GM Lloyd, AS Coomes, LN Eidson, ME De Sousa Rodrigues, DL Oliver, SD Kelly, J Chang, N Bengoa-Vergniory, R Wade-Martins, BI Giasson, V Joers, D Weinshenker, MG Tansey

## Abstract

Degeneration of locus coeruleus (LC) neurons and dysregulation of noradrenergic signaling are ubiquitous features of Parkinson’s disease (PD). The LC is among the first brain regions affected by α-synuclein (asyn) pathology, yet how asyn affects these neurons remains unclear. LC-derived norepinephrine (NE) can stimulate neuroprotective mechanisms and modulate immune cells, while dysregulation of NE neurotransmission may exacerbate disease progression, particularly non-motor symptoms, and contribute to the chronic neuroinflammation associated with PD pathology. Although transgenic mice overexpressing asyn have previously been developed, transgene expression is usually driven by pan-neuronal promoters and thus has not been selectively targeted to LC neurons. Here we report a novel transgenic mouse expressing human wild-type asyn under control of the noradrenergic-specific dopamine β-hydroxylase promoter. These mice developed oligomeric and conformation-specific asyn in LC neurons, alterations in hippocampal and LC microglial abundance, upregulated GFAP expression, degeneration of LC fibers, decreased striatal dopamine (DA) metabolism, and age-dependent behaviors reminiscent of non-motor symptoms of PD that were rescued by adrenergic receptor antagonists. These mice provide novel insights into how asyn pathology affects LC neurons and how central noradrenergic dysfunction may contribute to early PD pathophysiology.

**Significance statement:** α-synuclein (asyn) pathology and loss of neurons in the locus coeruleus (LC) are two of the most ubiquitous neuropathologic features of Parkinson’s disease (PD). Dysregulated NE neurotransmission is associated with the non-motor symptoms of PD including sleep disturbances, emotional changes such as anxiety and depression, and cognitive decline. Importantly, loss of central NE may contribute to the chronic inflammation in, and progression of, PD. We have generated a novel transgenic mouse expressing human asyn in LC neurons to investigate how increased asyn expression affects the function of the central noradrenergic transmission and associated behaviors. We report cytotoxic effects of oligomeric and conformation-specific asyn, astrogliosis, LC fiber degeneration, disruptions in striatal dopamine metabolism, and age-dependent alterations in non-motor behaviors without inclusions.

## Introduction

Brain regions affected in Parkinson’s disease (PD) exhibit proteinaceous aggregates (known as Lewy bodies) primarily composed of α-synuclein (asyn), chronic inflammation, and neuron loss (den Hartog and Bethlem, 1960; Spillantini et al., 1997; Tansey and Goldberg, 2010). In addition to deficiencies in striatal dopamine (DA) and the DA transporter (DAT), acetylcholine, serotonin, and norepinephrine (NE) systems are also affected (Gonera et al., 1997; Abbott et al., 2005; Ross et al., 2008).

The locus coeruleus (LC) is among the first brain regions affected in PD. The LC is a bilateral pontine nucleus at the lateral floor of the fourth ventricle, and is the main source of NE for the central nervous system (CNS) (Iversen et al., 1983; Mann et al., 1983; Braak et al., 2001). Asyn aggregates and neuronal degeneration in the LC are ubiquitous features of PD and are associated with non-motor symptoms including sleep disorders, mood disturbances, and cognitive deficits (Iversen et al., 1983; Chui et al., 1986; German et al., 1992; Braak et al., 2001; Zarow et al., 2003; Weinshenker, 2018). Imaging and histological studies show a progressive loss of central NE, noradrenergic neurons, and accumulation of asyn pathology in the LC in PD (Halliday et al., 1990; German et al., 1992; Brunnstrom et al., 2011; Pifl et al., 2012; Keren et al., 2015) which may exacerbate degeneration of DA neurons in the midbrain substantia nigra pars compacta (SNpc) (Zarow et al., 2003; Chen et al., 2014).

Depleting LC-NE exacerbates 6-hydroxydopamine (6-OHDA) or 1-methyl-4phenyl-1,2,3,6-tetrahydropyridine (MPTP)-induced nigrostriatal pathology in rodents and primates (Mavridis et al., 1991; Srinivasan and Schmidt, 2003; Rommelfanger et al., 2007), while increasing extracellular NE is protective (Kilbourn et al., 1998; Rommelfanger et al., 2004; Kreiner et al., 2019). Furthermore, lesioning LC neurons induces inflammation, and dysregulated NE neurotransmission may contribute to the chronic inflammation seen in PD (Kim and Joh, 2006; Tansey and Goldberg, 2010; Yao et al., 2015; Bharani et al., 2017; Song et al., 2018).

The initiating event in asyn aggregation in sporadic PD is unclear, but candidate mechanisms include increased expression of asyn, as individuals with a multiplication mutation in the gene encoding asyn (*SNCA*) develop autosomal dominantly-inherited PD (Singleton et al., 2003; Chartier-Harlin et al., 2004; Ferese et al., 2015). Age is the primary risk factor for PD, and rodent models of asyn overexpression develop age-dependent asyn aggregates and PD-like behavioral abnormalities (Masliah et al., 2000; Giasson et al., 2002; Hansen et al., 2013). However, in most of these models, transgene expression is driven by a pan-neuronal promoter with asyn overexpression in multiple regions of the CNS, with only one model reporting LC pathology (Masliah et al., 2000; Giasson et al., 2002; Maskri et al., 2004; Schell et al., 2009; Koprich et al., 2010; Sotiriou et al., 2010; Delenclos et al., 2017). Viral-mediated expression has been used to target asyn overexpression to specific brain regions (Baekelandt et al., 2002; Kirik et al., 2002; Delenclos et al., 2017; Ip et al., 2017; Niu et al., 2018). Notably, viral overexpression of a familial PD mutant asyn in LC neurons resulted in asyn aggregation, inflammation, and degeneration of LC neurons (Henrich et al., 2018).

To investigate specifically how pathology induced by increases in asyn affects noradrenergic neurons in the LC in an aging organism, we targeted expression of human wild-type asyn to LC neurons under control of the noradrenergic/adrenergic-specific dopamine β-hydroxylase (DBH) promoter using bacterial artificial chromosome (BAC) transgenesis. To determine the molecular, cellular, and behavioral age-dependent consequences of increased asyn expression in LC neurons, 3-, 14-, and 24-month (mo) old DBH-hSNCA transgenic (Tg) mice and non-transgenic (nTg) littermate controls were examined.

## Methods

### Experimental design

To determine how expression of human asyn in LC neurons affects neuronal health and LC-NE-associated behaviors, mice were aged undisturbed until the time of behavioral testing or sacrifice at ages 3-, 14-, and 24-mos.

### Generation of the DBH-hSNCA mouse model

Male and female mice expressing human wild-type α-synuclein (DBH-hSNCA) were engineered using a commercially available human bacterial artificial chromosomal (BAC) RP11-746P3 (Cubells et al., 2016) encompassing the *DBH* gene. The wild-type *hSNCA* cDNA open reading frame (400bp) was targeted to the translational start site of *DBH* by standard BAC recombineering methods by the University of North Carolina – Chapel Hill Molecular Neuroscience Core (currently Animal Model Core). The BAC construct was injected into C57BL/6N pronuclei by the Emory University Mouse Transgenic and Gene Targeting Core Facility (http://www.cores.emory.edu/tmc/index.html), transgene expression in founder pups was determined by PCR, and breeding lines were established. Mice carrying the hSNCA sequence were crossed with wild-type C57Bl/6N mice (Charles River) to establish the hemizygous transgenic DBH-hSNCA line.

To improve efficiency and accuracy of LC tissue isolation for western blot and mRNA analysis, DBH-hSNCA mice were crossed with the TH-EGFP reporter mouse expressing enhanced green fluorescent protein (EGFP) under the tyrosine hydroxylase (TH) promotor (Sawamoto et al., 2001).

### Animals

Male and female DBH-hSNCA mice were maintained on a C57Bl/6 background. Mice were group housed (maximum 5 mice per cage) until two weeks prior to the start of behavioral testing, when they were singly housed until euthanized. Animals were maintained on a 12/12h light/dark cycle with access to standard rodent chow and water *ad libitum.* Hemizygous animals served as experimental mice, with non-transgenic littermates as controls. Genotypes were determined by tail snip PCR with two sets of primers: Forward 5’ TGTCCAAGATGGACCAGACTC 3’ Reverse 3’ ACTGGTCTGAGGCAGGGAGCA 5’; Set Forward 5’ GCCCTCAGTCTACTTGCGGGA 3’ Reverse 3’ GCGAGAGCATCATAGGGAGT 5’. Experimental procedures involving use of animals were performed in accordance with the NIH Guidelines for Animal Care and Use and approved by the Institutional Animal Care and Use Committee at Emory University School of Medicine.

### Sleep latency test

Latency to fall asleep was quantified as the duration of time following gentle handling until their first sleep bout, which was defined as sleeping continuously for 2 min, and for a total of 75% of the 10-min period that began at sleep onset (Hunsley and Palmiter, 2004). Sleep testing began at 9 AM, 2 h into the light cycle when internal pressure to sleep is high. The sessions were video recorded and scored by an experienced observer blind to the genotype. We have validated this behavioral sleep scoring method with EEG (Porter-Stransky et al., 2019).

### Marble burying test

Marble burying was conducted as previously described (de Sousa Rodrigues et al., 2017) to determine whether expression of human asyn in LC neurons promotes anxiety-like behavior. Mice were placed in a plastic tub (50.5 x 39.4 x 19.7 cm) containing 5 inches of lightly pressed bedding. Twenty marbles of uniform size and color were placed in 5 rows of 4 marbles each on top of the bedding. Mice were placed in the containers and allowed to roam freely for 30 min. At the end of testing, the mice were placed back into home cages, and the number of marbles buried at least two-thirds of their height were counted. Marble burying was conducted 2 weeks after sleep latency testing.

### Open Field testing

In the open field test, a mouse that spends less time in or hesitates to re-enter the open center of the testing chamber is considered to be exhibiting anxiety-like behavior (Britton and Britton, 1981). During the light phase of the light/dark cycle, mice were acclimated to a dark testing room under red light for 1 h before testing. Mice were placed into the open field (45 cm X 45 cm square box) and allowed to move freely for 10 min. Distance, velocity, center, and border statistics were measured using Noldus/Ethovision software. Center was defined as the central 22.5 cm X 22.5 cm. Open field was conducted 1 week after marble burying.

### Circadian locomotion

All testing mice were acclimated to the testing room for 2 d prior to the experiment. Mice were each placed in a clear Plexiglas (15.75” L, 13.25” L, 7.38” H) activity cage equipped with infrared photobeams (San Diego Instruments, La Jolla, CA). Food and water were available *ad libitum* during the 23-h testing period. Ambulations (consecutive photobeam breaks) were recorded by PAS software. Circadian locomotion behavior was assessed 2 weeks after open field testing.

### Fear conditioning

Fear conditioning training and contextual and cued fear testing is a test of memory for the association of an aversive stimulus with an environment cue or context, and was conducted as previously described (Chalermpalanupap et al., 2018) over 3 consecutive days. Mice were placed in the fear conditioning apparatus (7” W, 7” D, 12” H, Coulbourn) with metal shock grid floor and allowed to explore the enclosure for 3 min. Following habituation, three conditioned stimulus (CS)-unconditioned stimulus (US) pairings were presented with a one-min inter-trial interval. The CS was a 20 sec 85 db tone, and the US was a 2 sec 0.5mA footshock (Precision Animal Shocker, Colbourn) which co-terminated with CS presentation. The contextual test was conducted on the following day when animals were placed back into the same chamber. On day three, the animals were placed in a novel compartment and allowed to habituate for 2 min. Following habituation, the 85 db tone was presented, and the amount of freezing behavior recorded. No shocks were given during the contextual or cued tests. Fear conditioning was conducted 1 week after circadian locomotion behavior.

### Behavioral pharmacology

For the three days prior to pharmacological experiments, mice received vehicle administration to habituate them to intraperitoneal (i.p.) injections. On the day of testing, mice received vehicle or a cocktail of the a1-adrenergic receptor antagonist prazosin (0.5 mg/kg; Sigma-Adrich, St. Louis, MO) and the β-adrenergic receptor antagonist DL-propranolol (5 mg/kg; Sigma-Aldrich), immediately prior to sleep latency testing. Drug doses were chosen based on previous studies and optimized in pilot experiments to ensure that behavioral effects were not due to sedation (Vazey and Aston-Jones, 2014). Treatments were counterbalanced between subjects, all mice received vehicle or drug cocktail with a seven-day washout period between testing days.

### Tissue collection

Animals used in immunohistochemical and high-performance liquid chromatography (HPLC) analyses were anesthetized by injection of sodium pentobarbital (Euthasol, Virbac) until unresponsive. Mice were transcardially perfused with phosphate-buffered saline (PBS; pH 7.4) until exiting blood ran clear. Brain tissue was removed, with one hemisphere post-fixed in 4% paraformaldehyde for immunohistochemistry, and the other dissected and flash frozen for HPLC. Animals used for qPCR or western blot analyses were euthanized by cervical dislocation under isoflurane anesthesia. Tissue was flash frozen and stored at −80°C until processing.

### Immunofluorescence and analysis

Brain tissue was sectioned on a freezing microtome (Leica SM2010R, Buffalo Grove, IL) at 40 μm and stored in cryoprotectant (30% ethylene glycol, 30% sucrose, 13.32 mM NaH_2_PO_4_, 38.74 mM Na_2_HPO_4_, 250 μM Polyvinylpyrrolidone) solution at −20°C until staining. Sections were washed in PBS before blocking in 5% normal goat serum (Jackson ImmunoResearch 005-000-121; NGS) with 0.05% Triton X-100 (Sigma #T9284100) in Tris-buffered saline pH 7.4 (TBS) for 1 h at room temperature. Sections were transferred directly to primary antibody solution containing 1% NGS, 0.05% Triton-X 100, and antibody at the concentrations described in Table 1 and incubated overnight at room temperature (05-02 Ms anti-NET) or 4°C (all other primary antibodies). Fluorescently conjugated secondary antibodies (described in Table 1) were diluted in 0.1% NGS with 0.05% Triton X-100, and tissue sections were incubated for 1 h at room temperature in the dark. Sections were mounted on Superfrost Plus slides (VWR) and were coverslipped with Vectashield with DAPI (Vector). All immunofluorescent images were acquired as z-stack images and the file compressed on a Keyence BZ-X700 microscope system (Itasca, IL). The Allen Brain Atlas version 1 (2008) was used to identify regions of interest (ROIs). One section per mouse containing the dorsal hippocampus (near bregma −1.995 mm) and one containing the LC (bregma - 5.555 mm) were analyzed for percent immunoreactivity (IR) within a standard ROI. A detection threshold was set uniformly across images in each analysis, and % IR determined using the “Measure” feature of ImageJ. Percent IR was calculated as area of IR within the ROI divided by the total ROI area and multiplied by 100. Quantification of Iba1-positive cells (microglia) was also analyzed with a standard threshold, ROI, and upper and lower size limits (pixel^2) using the “Analyze particles” function in ImageJ.

**Table 1:**
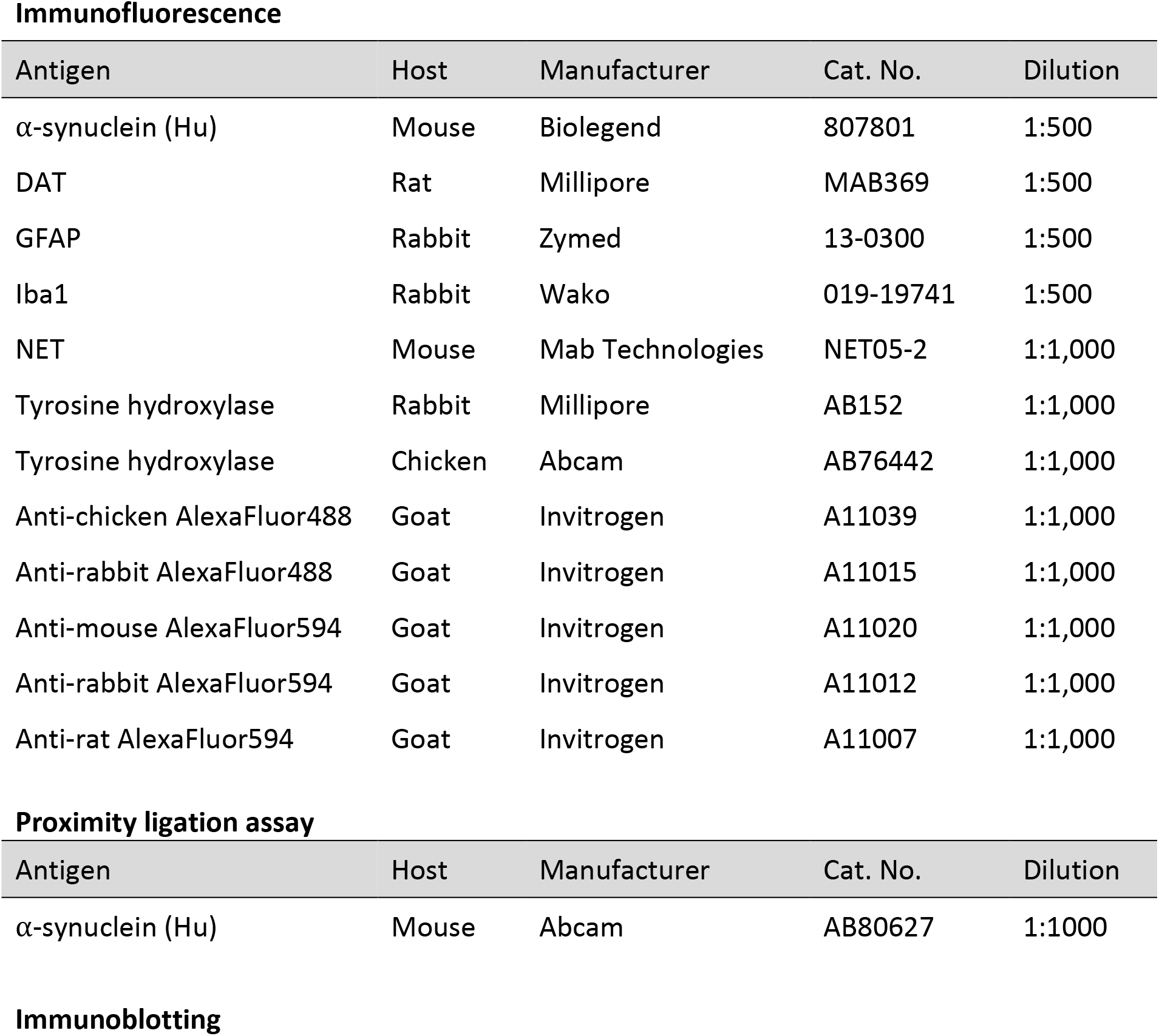
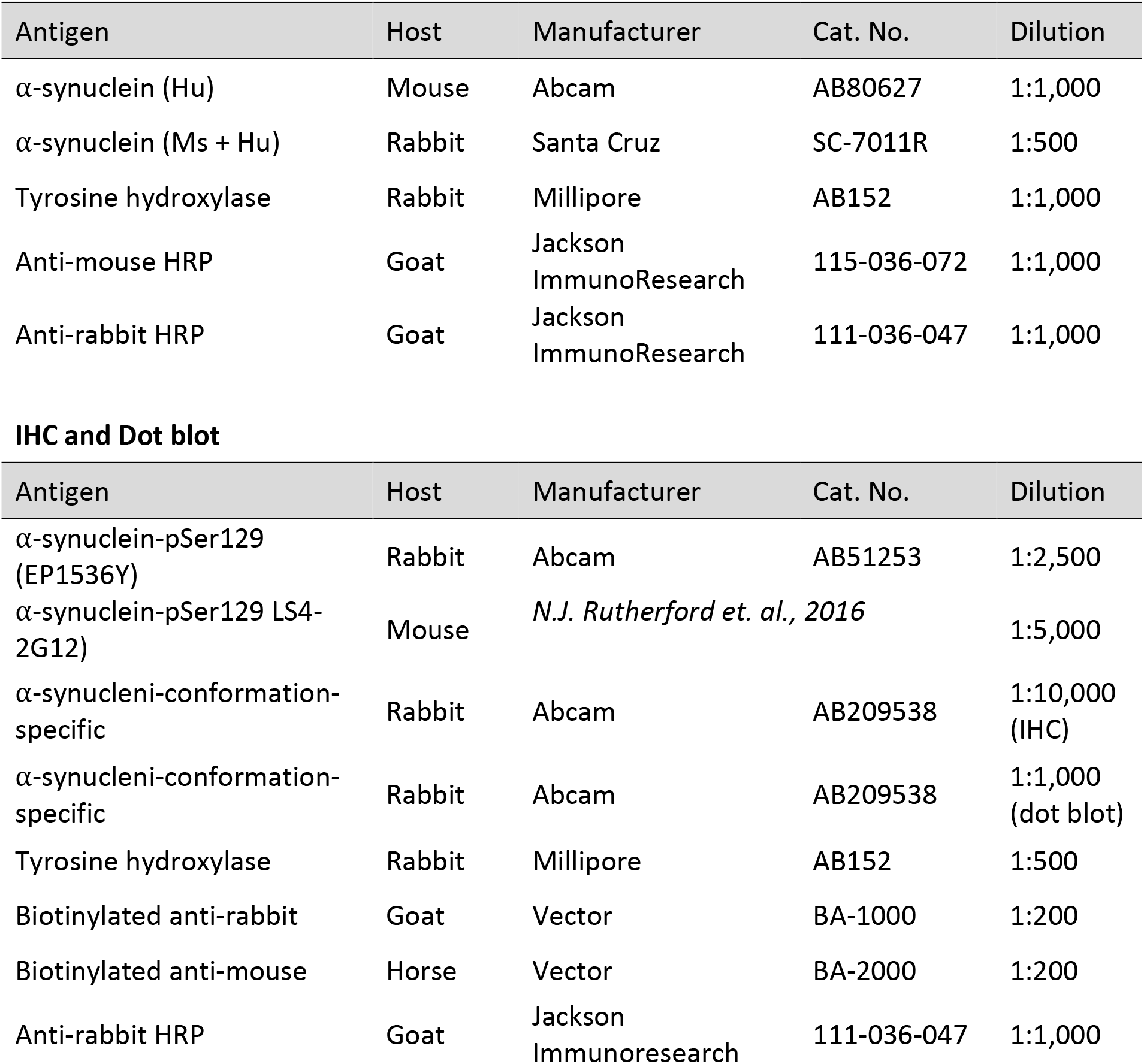
Primary and secondary antibodies used in these studies

### Immunohistochemistry and analysis

For immunoperoxidase visualization of pSer129 asyn (Rutherford et al., 2016) and conformation-specific asyn, brains were post-fixed with 4% paraformaldehyde, incubated in 70% ethanol, and then embedded in paraffin. Paraffin sections were cut at 5μm and stained using previously published protocols (Joers et al., 2014; Lloyd et al., 2020). Briefly, sections were deparaffinized and rehydrated, and heat-induced antigen retrieval was performed using 10mM sodium citrate solution (pH 6.0, pSer129) or citrate Target Retrieval Solution (Dako #S169984-2, pH 6.1) with 0.05% Tween-20 (conformation-specific) for 30 min. Next, endogenous peroxidase was blocked, and non-specific binding sites minimized with 8% serum (Jackson ImmunoResearch), 20% avidin/biotin blocking solution (Vector Laboratories #SP-2001), and 0.1% Triton-X 100 in TBS for 1 h (pser129) or with 2% FBS in 0.1M Tris (pH 7.6) for 10 min (conformation-specific). Sections were transferred into primary antibody solution containing 5% serum, and 20% avidin/biotin blocking solution (Vector Laboratories #SP-2001) in TBS (pSer129) or 2% FBS in 0.1M Tris (conformation-specific) and antibody concentrations detailed in Table 1 overnight at 4°C. The sections were then incubated in appropriate biotinylated secondary antibodies diluted with 5% serum in TBS (pser129) for 30 min or 2% FBS in 0.1M Tris for 1 h (conformation-specific) at room temperature. Antigen signal was amplified using VectaStain ABC-HRP Elite kit (Vector Laboratories #PK-6100) for 1 h at room temperature, and staining was developed using 3,3-diaminobenzine (DAB) as the chromogen (Sigma, #D4293). Sections were counterstained with Mayer’s hematoxylin (Sigma #51275). Sections stained for conformation-specific asyn underwent additional blocking with 2% FBS in 0.1M Tris before secondary antibody and DAB incubations. All DAB-immunostained slides were imaged with a Leica microscope (DMi8 Thunder) and post-processed through Adobe Photoshop to improve contrast. Intensity of conformation-specific asyn-IR in the LC were scored by experimenters blinded to genotype using the following rubric: 0 = no staining in the LC; 1 = < 25% of the LC stained; 2 = 25-50% of the LC stained; 3 = > 50% of the LC stained. Images for pser129 asyn were not quantified.

### Proximity ligation assay (PLA)

Paraffin-embedded tissue was rehydrated by consecutive incubations in xylene, Histoclear, 100% ethanol, 95% ethanol, 70% ethanol and H2O. Samples were then incubated in 10% H2O2 in PBS to reduce background and heated in a microwave in citrate buffer (pH 6.0; Abcam) for antigen retrieval. After antigen retrieval, samples were processed for immunofluorescence: 1 h RT incubation in 10% serum with 0.05% Tween-20 in TBS block, 1 h incubation in primary antibody, and TBS with 0.05% Tween-20 (TBS-T) wash. Slides were then incubated for 1 h with secondary antibodies conjugated to Alexa488 (Life Technologies) and washed again with TBS-T. For immunohistochemistry, samples were then coverslipped with FluorSave (Calbiochem). For PLA, samples were covered in manufacturers PLA blocking solution (Sigma) for 1 h at 37°C, and then incubated overnight with PLA conjugates (a-syn211; ab80627 Abcam). The next day, samples were washed with TBS-T, incubated in ligation solution for 1 h at 37° C, washed with TBS-T, incubated in amplification solution for 2.5 h at 37°C, washed with TBS, counterstained with DAPI, and mounted with FluorSave. All PLA reagents were used per manufacturer’s instructions (Sigma #92008). PLA puncta were counted by experimenters blinded to genotype from 25 cells per LC section per animal. Two independent experimental rounds were normalized and averaged to provide the final quantification.

### Dot blots

Frozen LC samples were homogenized in RIPA buffer (1% Triton-X 100, 50mM Tris HCL, 0.1% sodium dodecyl sulfate, 150mM NaCL, pH 8.0), or Trizol (Life Technologies #15596-018) and centrifuged at 12,000 x g for 10 min. Supernatants were transferred to fresh tubes and quantified for protein concentration by BCA assay. Samples were diluted in PBS, and 225 ng was spotted directly onto nitrocellulose membranes (Biorad #1620117) for dot blot analysis (Sampson et al., 2016). Membranes were air-dried for 10 min and blocked in TBS-T containing 5% non-fat dry milk for 1 h. Membranes were probed for conformation-specific asyn (Table 1) at 4°C overnight with gentle rocking. Membranes were incubated with HRP-conjugated secondary antibodies, followed by SuperSignal™ West Pico PLUS Chemiluminescent Substrate (ThermoScientific) and imaged using a Licor (Odyssey FC) system.

### RNA Scope

*In situ* RNA analysis was performed using RNAScope Multiplex Fluorescent v2 kit (ACD Bio 3231000). Tissue prep and analysis were conducted as described in manufacturer’s protocol. Briefly, following transcardial perfusion with saline, brains were incubated in 4% paraformaldehyde (PFA) for 24 hours followed by a series of increasing sucrose concentrations before being frozen in optimal temperature cutting medium (Sakura) and stored at −80°C until sectioning. Tissue sections (12μm) were collected on a Leica CM1900 cryostat and mounted on Superfrost Plus slides. To prevent tissue detachment, slides were dried at 60°C for 30 min and fixed in 4% PFA for 15 min at 4°C before ethanol dehydration. Tissue was processed as described in ACD Bio protocol (acdbio.com/technical-support/user-manuals #323100-USM).

### Western immunoblotting

Western blots were conducted as previously described (de Sousa Rodrigues et al., 2017). Flash frozen samples were stored at −80°C until processing. Protein was isolated from LC samples with RIPA buffer. RIPA samples were centrifuged at 12,000 rpm for 20 min at 4°C. Supernatant was transferred to new tube for bicinchoninic acid protein assay (Pierce Scientific #23225). Trizol samples were resuspended in 1% SDS. Samples were diluted to 1μg/μl in 4x sample buffer (BioRad #1610747) and boiled at 90°C for 5 min. Electrophoresis was performed using 12% gels (BioRad #4568046; 5μl) and transferred to 0.45 μm PVDF membrane using Trans-Blot Turbo Transfer System (BioRad). For immunoblotting, the membrane was fixed in 0.4% PFA for 30 min following transfer. After a brief wash, blots were incubated in 5% milk blocking buffer (BioRad) for 1 hour at 4°C before primary antibody overnight at 4°C (See Table 1). Membranes were washed with TBS-T (0.01% Tween-20) and incubated in HRP-conjugated secondary antibodies in blocking buffer for 1 hour at room temperature. Images were acquired using Azure Biosystems and analyzed by ImageStudio Lite software. Protein expression was normalized to total protein (Li-Cor #926-11015).

### High performance liquid chromatography (HPLC)

Monoamines were examined by high performance liquid chromatography with electrochemical detection as described previously (Song et al., 2012). For HPLC, an ESA 5600A CoulArray detection system, equipped with an ESA Model 584 pump and an ESA 542 refrigerated autosampler was used. Separations were performed using an MD-150 × 3.2 mm C18 (3 μM) column at 25°C. The mobile phase consisted of 8% acetonitrile, 75 mM NaH2PO4, 1.7 mM 1-octanesulfonic acid sodium and 0.025% trimethylamine at pH 2.9. Twenty-five microliters of sample were injected. The samples were eluted isocratically at 0.4 mL/min and detected using a 6210-electrochemical cell (ESA, Bedford, MA) equipped with 5020 guard cell. Guard cell potential was set at 475 mV, while analytical cell potentials were −175, 150, 350 and 425 mV. The analytes were identified by the matching criteria of retention time and sensor ratio measures to known standards (Sigma Chemical Co., St. Louis MO.) consisting of dopamine, norepinephrine, 3,4-dihydroxyphenylacetic acid (DOPAC), and 4-Hydroxy-3-methoxyphenylglycol (MHPG). Compounds were quantified by comparing peak areas to those of standards on the dominant sensor.

### Statistical analysis

Student’s t-test was used to assess differences by genotype within each age group in sleep latency, marble burying, open field, western blot, and all immunofluorescent analyses. Repeated measures two-way ANOVA was used to analyze differences by genotype (between-subject) for within-subject conditions including fear conditioning and circadian locomotor assay followed by Tukey’s post hoc test where applicable. Analyses were conducted within each age group. Comparisons across age groups were not conducted, as behavioral assays, HPLC, and immunofluorescence of each cohort were conducted at separate time points. The analyses were performed using GraphPad Prism 7 (GraphPad Software, Inc., La Jolla, CA) with a p-value threshold of <0.05.

## Results

### Generation of DBH-hSNCA mice

The DBH-hSNCA mouse model was developed using a DBH-BAC construct carrying the wild type human *SNCA* cDNA open reading frame at the translational start site of *DBH* (Fig. 1). Transgene integration was confirmed by PCR, and founder mice were bred with wild type C57BL/6 mice to establish the hemizygous DBH-hSNCA line.

**Figure 1:**
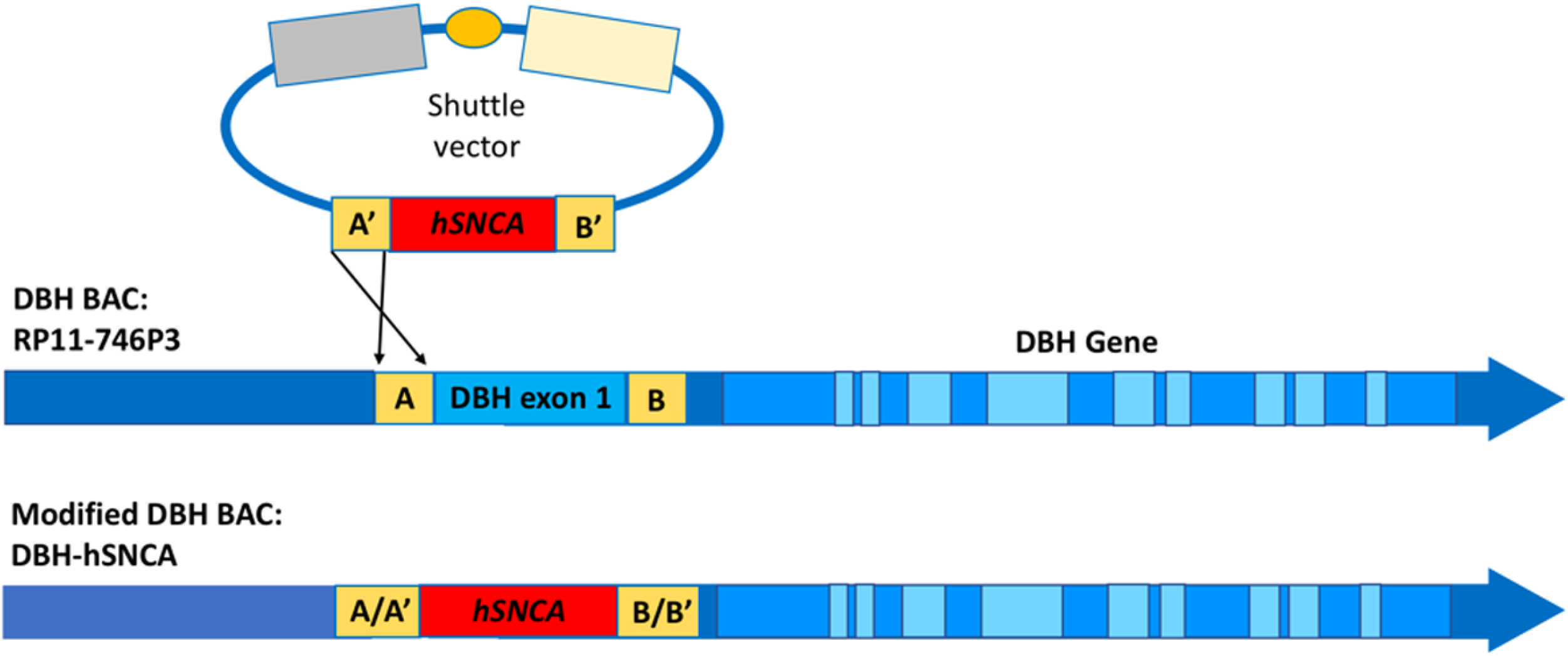
*hSNCA* cDNA open reading frame was targeted to the translational start site of *DBH* by standard BAC recombineering methods

### Human *Snca* mRNA is expressed in DBH-hSNCA LC neurons

Fluorescent *in situ* mRNA analysis revealed human *Snca* mRNA in Tg LC neurons (Fig. 2), which co-localized with mouse *Th* and mouse *Dbh* mRNA, while human *Snca* mRNA expression was not detected in nTg LC neurons.

**Figure 2:**
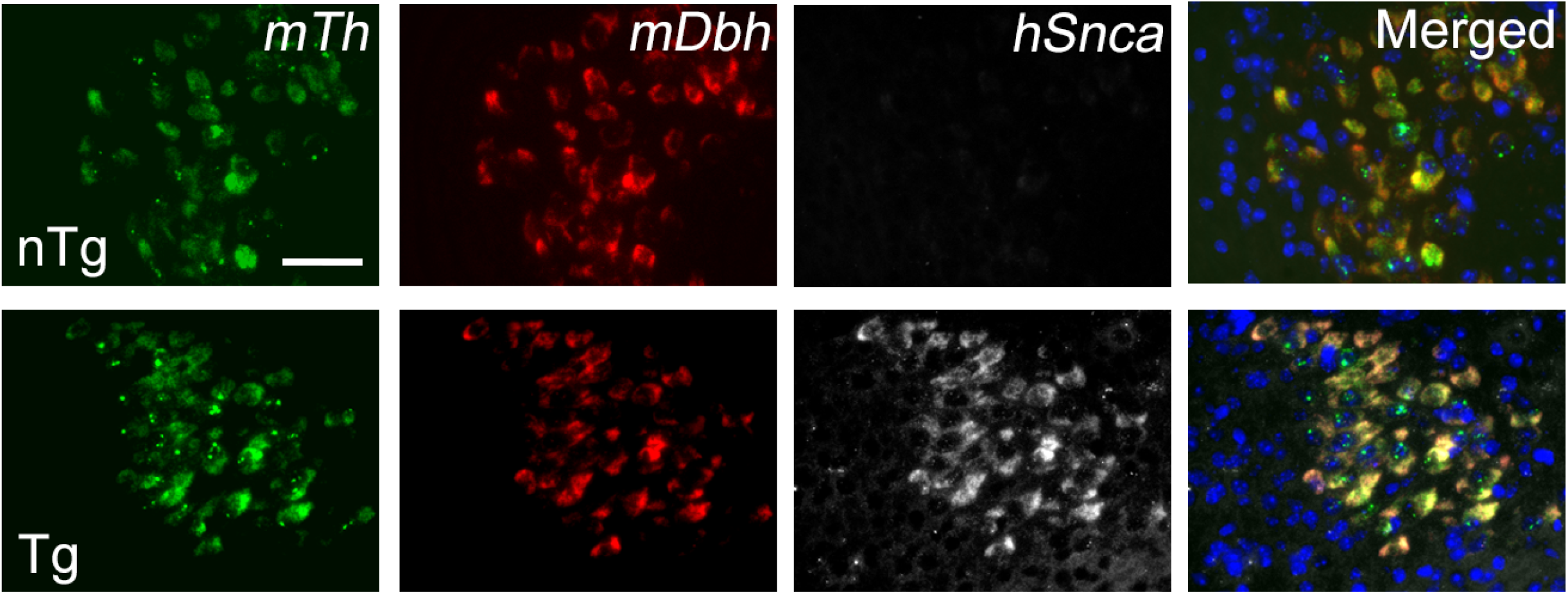
Human *Snca* mRNA is expressed in DBH-hSNCA locus coeruleus neurons. Human *Snca* mRNA (hSnca; white) expression is detectable only in transgenic (Tg: bottom row) LC neurons, where it co-localizes with mouse *Th* mRNA (mTh, green), and mouse *Dbh* mRNA (mDbh; red) using RNA Scope Fluorescent Multiplex v2 assay. Scale bar = 50μm.

### Human asyn is expressed in select noradrenergic brainstem nuclei of DBH-hSNCA transgenic mice

Human asyn protein in the LC was analyzed by immunofluorescence and western blot. Using an antibody specific for human asyn (Biolegend 807801), expression of human asyn was found to co-localize with TH-expressing LC neurons only in brain sections from DBH-hSNCA Tg mice (Fig. 3A). No human asyn-specific immunofluorescence was detected in LC neurons of nTg littermates or in SNpc neurons regardless of genotype (Fig. 3A,C). Human asyn was also detectable specifically in LC protein lysate from Tg tissue by immunoblot and not in nTg lysates (Fig. 3B; n=4). Quantitative western blot analysis of LC protein using a pan-asyn antibody to detect both human and mouse asyn protein revealed a significant ~30% increase of total asyn in Tg mice at 3-mo relative to that in nTg littermates (Fig. 3D; t_(6)_=3.156, p=0.0197, n=4). Human *SNCA* mRNA expression in Tg LC neurons was confirmed by qPCR (Data not shown). Expression of human asyn was confirmed in the A4 (Fig. 3G) and A5 (Fig. 3H) DBH-expressing regions in the pons of Tg mice by immunofluorescence, but expression was not detectable in the medullary A1/C1 (Fig. 3E) or A2/C2 (Fig. 3F) nuclei of DBH-hSNCA mice.

**Figure 3:**
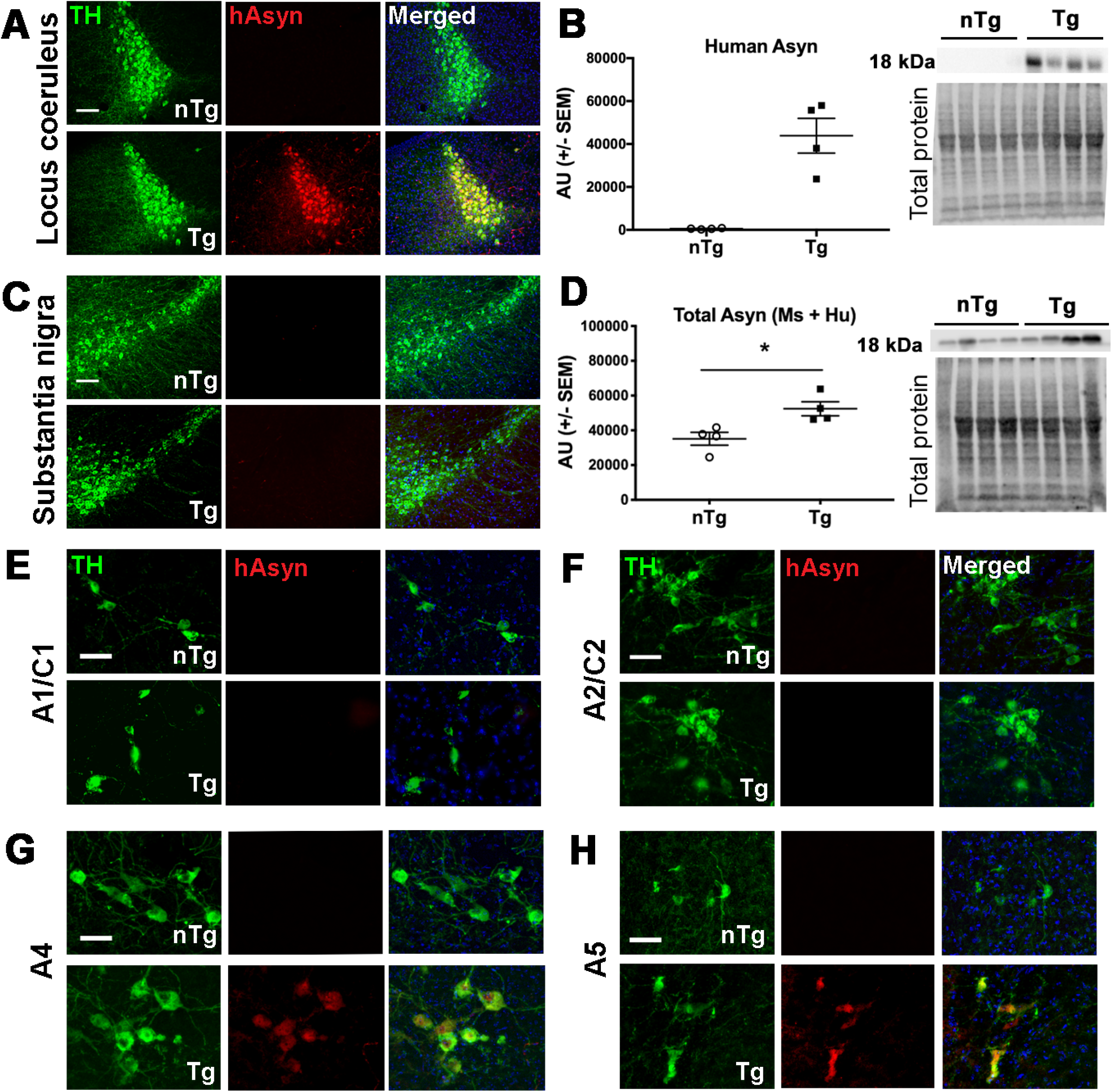
Analysis of asyn expression in catecholaminergic brainstem regions of young DBH-hSNCA mice. ***A,*** Immunofluorescent detection of human asyn (red) with a species-specific antibody (Biolegend 807801) demonstrates co-localization with TH-expressing LC neurons (green) in Tg mice (lower panel) but not in nTg mice (upper panel), or in ***C,*** TH-expressing neurons (green) in the substantia nigra regardless of genotype. ***B,*** Human asyn protein is expressed selectively in LC neurons of Tg mice by western blot. ***D,*** Immunoblot of LC protein with an antibody against asyn that detects the mouse and human protein reveals a significant increase in total asyn protein expression in Tg LC neurons as compared to that in LC of nTg littermate mice. Immunoblot data graphed as arbitrary units (AU) normalized to total protein. Human asyn is not expressed in the ***E,*** A1/C1 or ***F,*** the A2/C2 region, but is detectable in the ***G,*** A4 and ***H,*** A5 regions of Tg mice by immunofluorescence. All data are from 3-mo old nTg and Tg mice. Scale bar = 100μm (A, C) or 50 μm (E-H). Student’s t-test ± SEM *\*p*<0.05.

### DBH-hSNCA mice exhibit behavioral phenotypes that resemble features of non-motor PD symptoms

A primary role of the LC-NE system is promoting arousal and wakefulness; LC activity is highest just prior to, and during wake (Hobson et al., 1975). Sleep disturbances are one of the most common non-motor PD symptoms, and PD patients with disturbed sleep have greater asyn pathology in the LC than PD patients without sleep complaints (Kalaitzakis et al., 2013). Thus, sleep latency was assessed to examine whether features of the sleep/wake cycle were affected by human asyn expression in the LC. Mice were gently handled and returned to their home cage, and video recording was scored by an observer blind to the genotype to determine latency to fall asleep. Our findings indicate that there was a significant increase in sleep latency in Tg mice at 3-mos, (Fig. 4A; t_(14)_=4.36 p=0.0007; n=8) and 14-mos (t_(17)_=2.51, p=0.0225; n= 9 nTg & 10 Tg), indicative of an elevated arousal state. No differences were observed at 24-mos (t_(8)_=0.821, p=0.4354, n= 8 nTg & 7 Tg).

**Figure 4:**
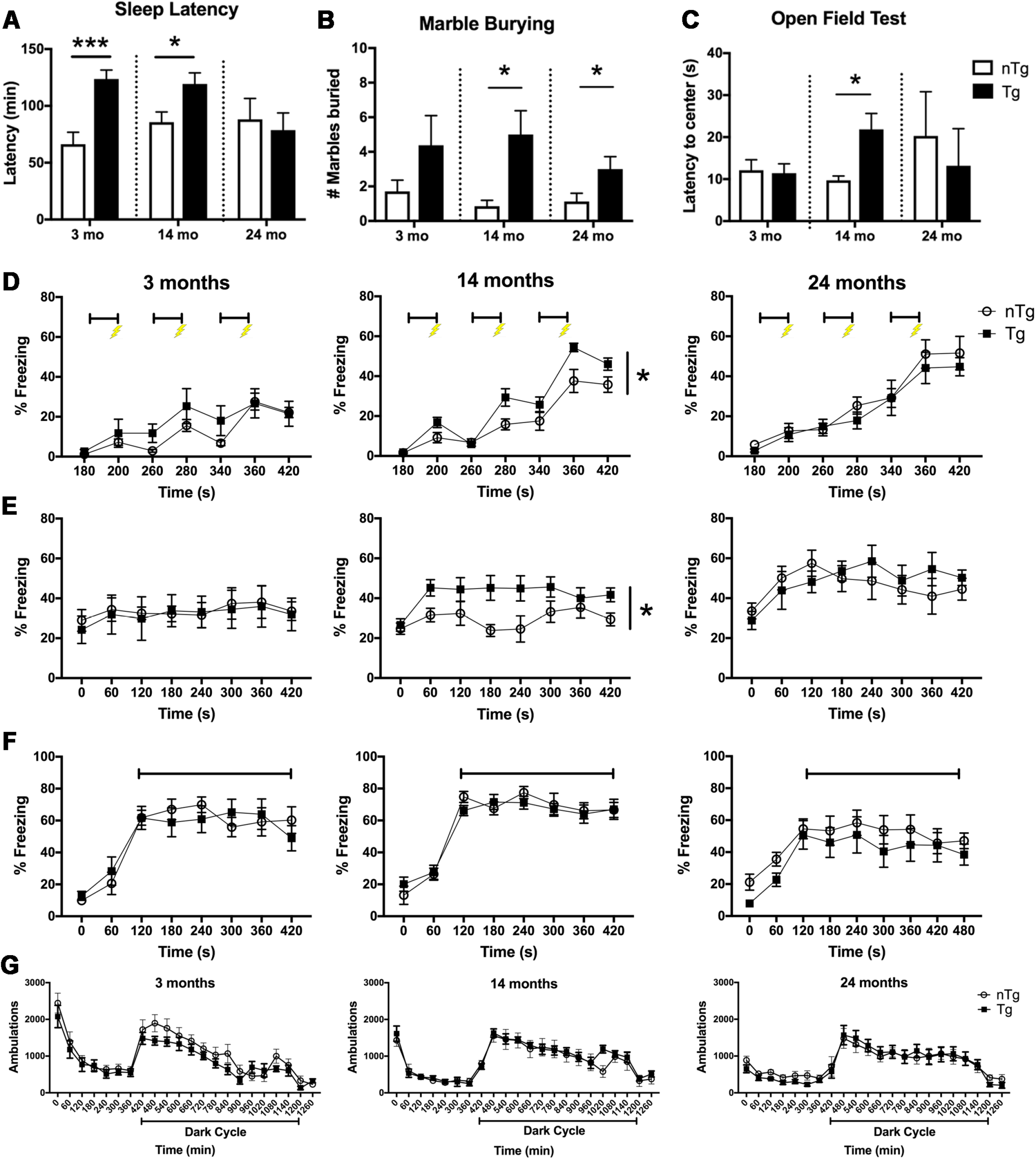
DBH-hSNCA mice display age-dependent behavioral abnormalities similar to the non-motor symptoms of PD. ***A,*** 3-mo and 14-mo Tg mice have increased sleep latencies. ***B,*** Tg mice display anxiety-like behavior by marble-burying test at 14-mos and 24-mos and ***C,*** latency to re-enter the center in the open field test at 14-mos. ***D,*** There is a significant effect of genotype at 14-mos with Tg mice exhibiting significantly more percent time freezing during fear training (center panel; bars represent tone presentation, bolt represents foot shock) and ***E,*** contextual learning. ***F,*** There was no effect of genotype in percent time freezing during cued test at any age. ***G,*** Ambulatory behavior does not differ by genotype at any age. Students t-test of genotype ± SEM in A, B & C. Repeated measures ANOVA ± SEM in D, E, F, and G. *\*p*<0.05, *\*\*\*p*<0.001.

Central noradrenergic neurons, including the LC and A2 regions, are activated by stress exposure, and blocking LC activity abolishes stress-induced anxiety-like behavior (Rinaman, 2011; McCall et al., 2015). In PD, anxiety severity is inversely correlated with LC function (Remy et al., 2005). To examine anxiety-like behavior, the behavior DBH-hSNCA mice was assessed in marble burying and open field tests. In the marble burying assay, there was a significant increase in the number of marbles buried by Tg mice at 14-(Fig. 4B; t_(18)_=2.735, p=0.0136; n= 8 nTg & 12 Tg) and 24-mos (t_(13)_=2.212, p=0.0455, n= 8 nTg & 7 Tg) but not at 3-mos (t_(13)_=1.369, p=0.1942, n= 7 nTg & 8 Tg). In open field tests, an age-dependent anxiety-like phenotype was also evident in 14-mo Tg mice, evinced by an increase in the latency to re-enter the center of the testing field (Fig. 4C; t_(17)_2.359, p=0.0305; n= 7 nTg & 12 Tg), with no differences in 3-(t_(11)_=0.1988, p=0.8460, n= 8 nTg & 5 Tg) or 24-mo old animals (t_(11)_=0.5075, p=0.6219, n= 5 nTg & 5 Tg). The latency to enter the center score for one 24-mo nTg animal was omitted after failing to enter the center during the open field test.

Fear conditioning is a measure of hippocampal-dependent (contextual) or -independent (cued) associative learning (Phillips and LeDoux, 1992), and mice lacking NE exhibit impaired contextual learning (Murchison et al., 2004). In a standard fear conditioning paradigm, 14-mo old DBH-hSNCA Tg mice exhibited increased freezing behavior during the fear training session (Fig. 4D center; Interaction F_(6, 120)_=2.735, p=0.0159; n=8-14), as well as during the contextual test (Fig. 4E center; effect of genotype F_(1, 20)_=5.566, p=0.0286), with no differences in freezing behavior during the cued test (Fig. 4F; Interaction F_(7, 160)_=0.5146, p=0.8128). No genotype differences were found at 3- or 24-months in the fear training (Fig. 4D; 3-mos Interaction F_(6,72)_= 1.082, p=0.3815, n=7; 24-mos Interaction F_(6, 78)_=0.3783, p=0.8908, n = 8 nTg & 7 Tg), contextual test (Fig. 4E; 3-mos Interaction F_(7, 84)_= 0.1477, n=7; p=0.9938; 24-mos Interaction F_(7, 91)_=1.040, p=0.4089, n = 8 nTg & 7 Tg) or cued test (Fig. 4F; 3-mos Interaction F_(8, 96)_=1.508, p=0.1645, n=7; 24-mos Interaction F_(8, 104)_=0.2766, p=0.9723, n = 8 nTg & 7 Tg).

Because disruption in locomotor activity can affect behavioral testing, we evaluated the locomotion over 24 h and found no genotype differences at 3-mos (Interaction F_(21, 252)_=1.033, p=0.4233; n=7), 14-mos (Interaction F_(21, 231)_=0.6319, p=0.8929; n= 7 nTg & 6 Tg), or 24-mos (F_(21, 273)_=0.06975, p=0.8349; n= 8 nTg & 7 Tg) of age (Fig. 4F). Initially, all groups had high levels of activity, as would be expected in a novel environment, which decreased as mice habituated to the test apparatus. Ambulations increased normally in all genotypes at commencement of the dark phase, when mice are typically more active, and decreased once the next light cycle began.

### Sleep latency in 3-, and 14-mo old DBH-hSNCA mice is normalized by adrenergic receptor blockade

The behavioral results above (i.e. increased arousal and anxiety-like behavior) were consistent with overactive LC-NE transmission. To test this hypothesis, an additional cohort of 3- and 14-mo old DBH-hSNCA mice underwent sleep latency testing following administration of saline or a cocktail of propranolol (β-adrenergic receptor antagonist) and prazosin (α1-adrenergic receptor antagonist). Similar to baseline testing, 3- and 14-mo Tg mice administered saline took longer to fall asleep than nTg littermates. However, adrenergic receptor blockade normalized sleep latency in 3-mo old or 14-mo old Tg mice, while propranolol and prazosin administration had no effect on sleep latency in nTg mice (Fig. 5A; Interaction F_(1, 16)_=7.367, p=0.0153, n=5) (Fig. 5B; Interaction F_(1, 18)_=1.27, p=0.0035, n= 6 nTg & 5 Tg).

**Figure 5:**
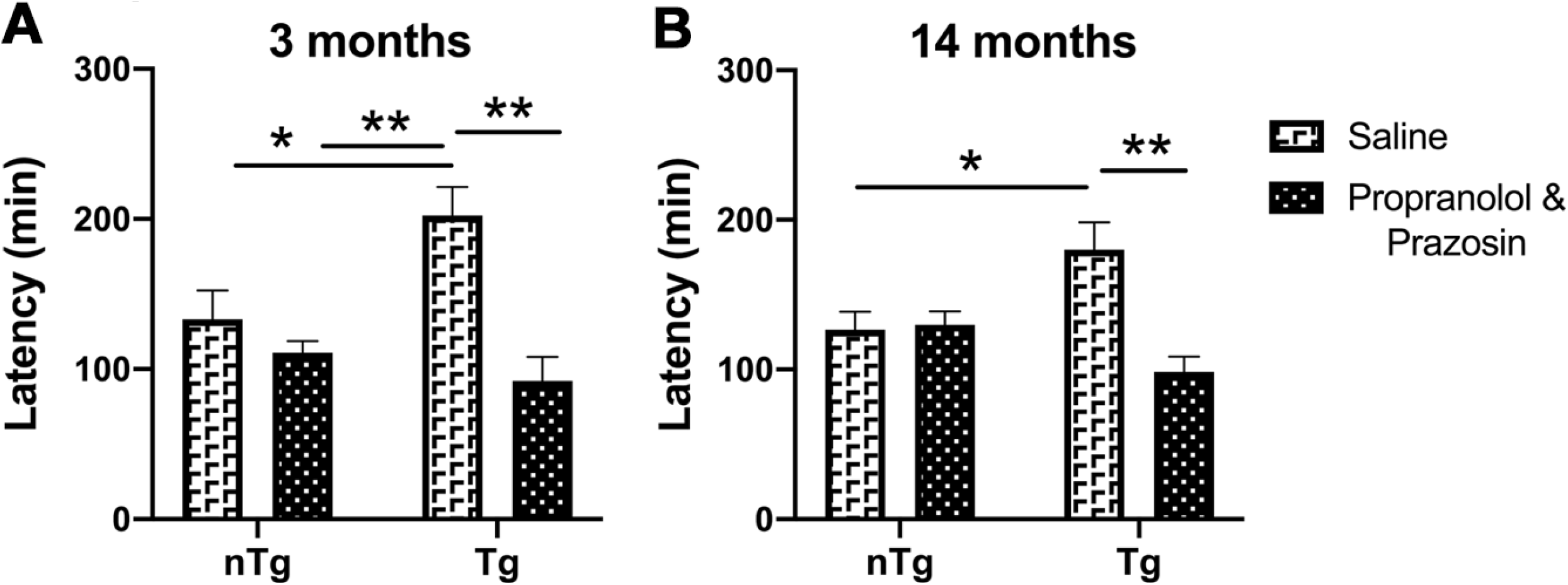
Adrenergic receptor antagonists normalize sleep latency in 3- and 14-mo old DBH-hSNCA mice. ***A,*** The increased sleep latency observed in 3-mo, and ***B,*** 14-mo old Tg mice was reduced by a cocktail of propranolol (β-adrenergic receptor antagonist) and prazosin (α1-adrenergic receptor antagonist), while no effect was observed in nTg mice at either age. Two-way ANOVA ± SEM. \**p*<0.05, \*\**p*<0.01.

### Human asyn forms oligomers and conformation-specific asyn in LC neurons at 14- and 24-mos of age

To determine whether transgenic expression of human asyn in LC neurons resulted in formation of asyn oligomers, tissue sections containing the LC were analyzed using a human asyn proximity ligation assay (PLA). Tg LC neurons displayed more asyn puncta per TH-positive neuron than nTg LC neurons at 14 (t_(8)_=2.53, p=0.0352; n=4 nTg & 6 Tg) and 24 months of age (t_(8)_=2.402, p=0.0430; n=12 nTg & 3 Tg) (Fig. 6A,B). To investigate whether the oligomeric asyn had altered conformation, we probed LC tissue lysates from 3-month old Tg and nTg mice for immunoreactivity against an asyn filament conformation-specific antibody (MJFR14) and tyrosine hydroxylase (TH) on immune-dot blots, using Snca knock-out and M83 asyn transgenic mouse tissues as negative and positive controls, respectively. We found that only Tg mice displayed robust MJFR14-IR (Fig. 6C), and it trended to be negatively correlated to TH in the LC (Fig. 6D), suggesting increased asyn may downregulate TH expression and/or induce dysfunction. We next probed for conformation-specific asyn properties using an immunohistological approach and found significant variability in the intensity of MJFR14-IR in LC neuron sections of 24-month old Tg and nTg mice that did not reach statistical significance between genotypes (Fig. 6E,F). Lastly, because conformation-specific and aggregated asyn has been associated with increased phosphorylation at serine 129 (pSer129), and labeling these forms of the protein is commonly used to identify asyn inclusions (Fujiwara et al., 2002; Wakamatsu et al., 2007; Schell et al., 2009), we immunolabeled LC tissue sections with antibodies specific for fibril-rich asyn (Syn-F1, data not shown) and asyn pSer129 (EP1536Y and LS4-2G12; Fig. 6G) and found no differences between genotypes, suggesting that at 24-mos of age there are no mature higher-order asyn inclusions in DBH-hSNCA mice.

**Figure 6:**
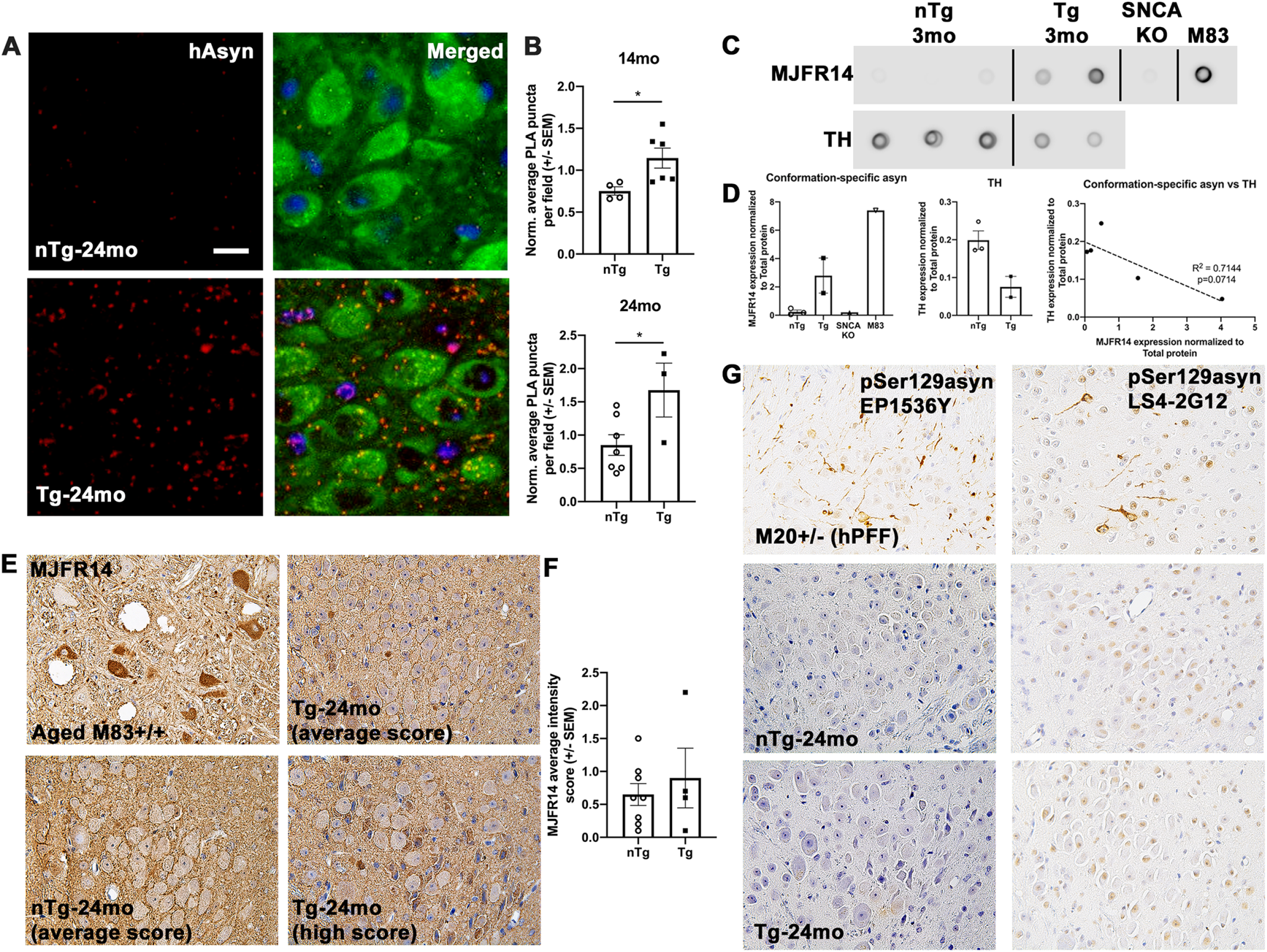
DBH-hSNCA LC neurons contain oligomeric and conformation-specific asyn. ***A,*** Representative images of oligomerized asyn (red) in LC neurons of (TH; green) 24-mos Tg mice as determined by proximity ligation assay (PLA). ***B***, Quantification of PLA puncta demonstrated significantly more oligomerized asyn (red) in LC neurons of Tg mice. ***C,*** Dot blot analysis of conformation-specific asyn and TH in the LC from 3-mo old nTg and Tg mice and in brain tissue from SNCA knock-out and M83^+/+^ asyn transgenic mice. ***D*** Quantification of MJFR14 and TH expression normalized to total protein from dot blots. Conformation-specific asyn trended to negatively correlate to TH in the LC of nTg and Tg mice. ***E,*** Representative microphotographs of DAB-immunostained LC sections from 24-mo old mice and positive control tissue from an aged M83^+/+^ asyn transgenic mouse with conformation-specific asyn (MJFR14). ***F,*** Semi-quantification analysis of MJFR14 intensity. ***G,*** Representative DAB-immunostained LC sections from 24-mo old mice and hippocampal positive control tissue from 6-mo old M20^+/-^ asyn transgenic mice after brain inoculation with human asyn pre-formed fibrils (hPFF) to seed asyn inclusions with antibodies against pSer129 asyn (EP1536Y and LS4-2G12). Scale bar (hAsyn) = 15μm. Scale bar (pser129 and MJFR14) = 100 μm. Student’s t-test ± SEM. *\*p*<0.05.

### Human asyn expression in LC neurons impacts striatal dopamine metabolism in 24-mo old mice

Dysregulated catecholamine metabolism and degeneration of catecholaminergic neurons are well-established features of PD (Iversen et al., 1983; Mann et al., 1983; Hirsch et al., 1988; Fearnley and Lees, 1991). Therefore, we measured catecholamine levels using high-performance liquid chromatography (HPLC). Hippocampal and striatal tissue content of NE, the NE metabolite MHPG, DA, and the DA metabolite DOPAC were quantified, revealing that NE (Fig. 7A,B) and DA (Fig. 7E,F) were not significantly affected in the hippocampus or striatum at any age. Similarly, the ratio of the major NE metabolite MHPG to NE was unaffected (Fig. 7C,D). The ratio of the DA metabolite DOPAC to DA in the hippocampus was unaltered (Fig. 7G), but was significantly reduced in the Tg striatum at 24-mos (Fig. 7H; t_(10)_=3.546, p=0.0046; n= 9 nTg & 6 Tg), consistent with decreased DA turnover.

**Figure 7:**
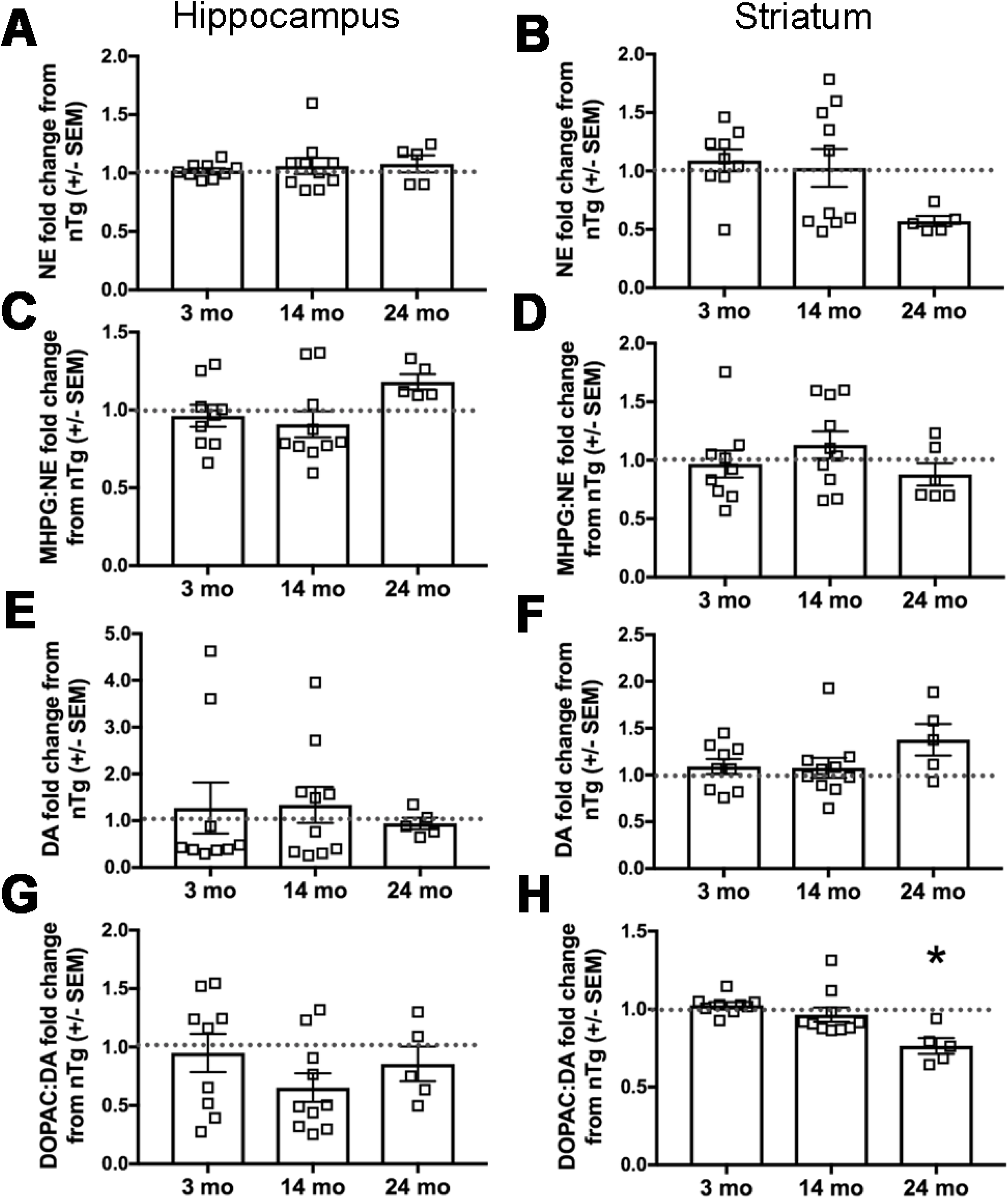
Human asyn expression in LC neurons impacts striatal dopamine metabolism in aged DBH-hSNCA mice. Catecholamine and catecholamine metabolite tissue content from hippocampus and striatum was measured by HPLC. ***A,*** Hippocampal NE is not affected by genotype. ***B,*** Striatal NE is reduced at 24-mos but does not reach statistical significance. The NE metabolite MHPG to NE ratio does not differ at any age in the ***C,*** hippocampus, or ***D,*** striatum. Dopamine content is not affected at any age in ***E,*** hippocampus, or ***F,*** striatum. ***G,*** The ratio of the DA metabolite DOPAC to DA is unaffected in the hippocampus ***H,*** but is significantly reduced in the striatum of Tg mice at 24-mos. Student’s t-test of genotype for each age group ± SEM. *\*p*<0.05.

### Human asyn expression does not affect LC neuronal integrity

LC neurons were visualized using NET-IR. NET is a reliable marker of LC neurons, and its expression is reduced in PD patients (Remy et al., 2005). Using a standard ROI, no difference in the percent NET IR was detected between genotypes at 3-(Fig. 8; t_(17)_=0.4537, p=0.6558; n= 9 nTg & 10 Tg), 14-(t_(10)_=0.8908, p=0.3939; n= 7 nTg & 5 Tg), or 24-mos (t_(8)_=0.8069, p=0.4430; n=5).

**Figure 8:**
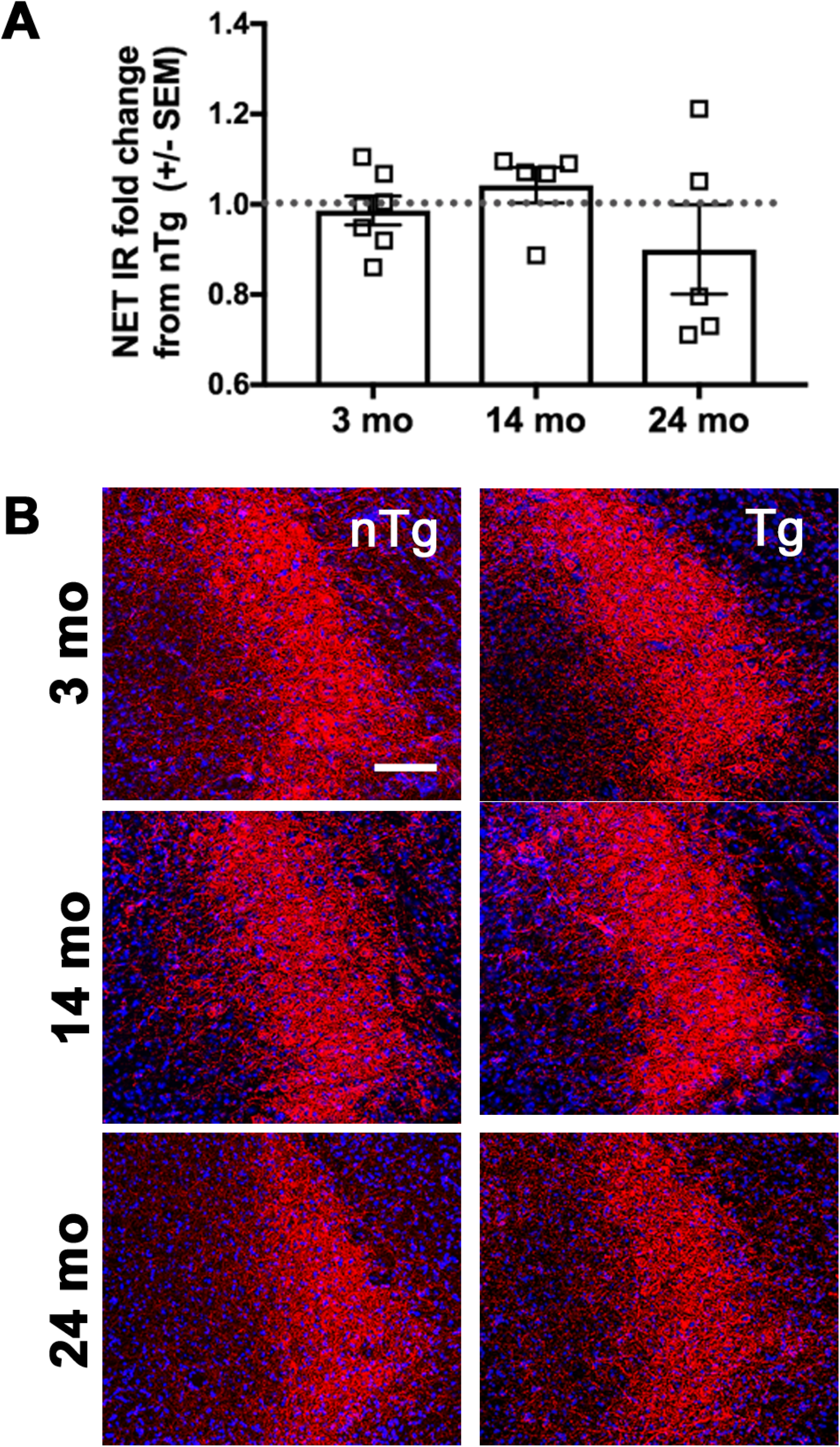
Human asyn expression does not affect LC integrity. NET-expressing LC cell bodies (red) percent immunoreactivity (IR) does not differ by genotype at 3-, 14-, or 24-mos. Student’s t-test of mean IR by genotype for each age group, ***A,*** graphed as fold change Tg from nTg mean ± SEM. ***B,*** Representative images. Scale bar = 50μm, *\*p*<0.05.

### Elevation of tyrosine hydroxylase in the LC of DBH-hSNCA mice

Tyrosine hydroxylase (TH) is the rate-limiting enzyme in NE and DA synthesis, and its long-term activity depends on its expression levels (Levitt et al., 1965; Haycock, 1993; Kumer and Vrana, 1996). To determine whether TH expression in the LC is affected by human asyn, we assessed TH IR in sections from 3-, 14-, and 24-mo old mice. TH IR was normalized to NET IR to control for potential differences in Bregma level between sections. TH expression was increased in Tg LC neurons (Fig. 9A,B) at both 3-(t_(17)_=2.154, p=0.0459; n= 9 nTg & 10 Tg) and 14-mo (t_(10)_=2.463, p=0.0335; n= 7 nTg & 5 Tg) of age. Western blot analysis from 3-mo old TH-EGFP-expressing LC neurons (Fig. 9C,D) confirmed higher TH expression in Tg animals (t_(10)_=3.837, p=0.0033; n= 6 nTg & 5 Tg).

**Figure 9:**
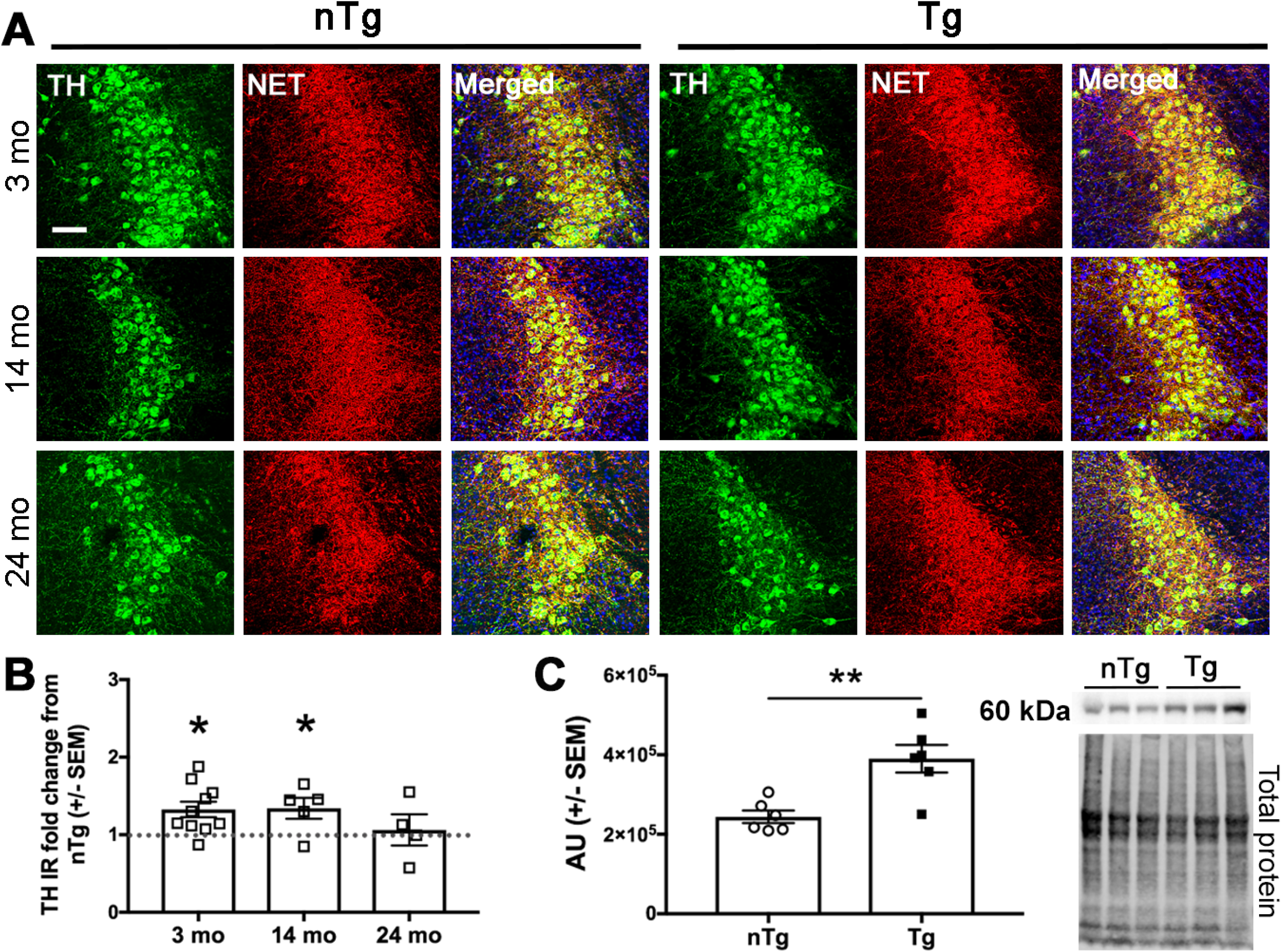
Tyrosine hydroxylase levels are elevated in young DBH-hSNCA Tg mice. Tyrosine hydroxylase (TH: green) immunoreactivity is increased in LC neurons (NET: red) at 3- and 14-mos of age. ***A,*** Representative immunofluorescent images. ***B***, TH IR mean normalized to NET IR mean and expressed as fold change Tg from nTg mean % IR ± SEM. ***C,*** Western blot analysis of 3-mo old LC neurons confirms increased TH abundance. Data graphed as arbitrary densitometric units (AU) was normalized to total protein ± SEM. ***D,*** Representative western blot images of TH (top) and total protein stain (bottom). Student’s t-test of nTg and Tg at each age, n=5-10. Scale bar = 50μm. *\*p*<0.05.

### Human asyn expression in LC neurons affects expression of local inflammatory markers

A wealth of studies suggest that dysregulated noradrenergic neurotransmission is associated with inflammation (reviewed by Butkovich et al., 2018), and that dysregulation of the LC-NE system could contribute to the chronic neuroinflammation observed in PD (Fujita et al., 1998; Gyoneva and Traynelis, 2013; Johnson et al., 2013). To determine whether human asyn expression in LC neurons affects the number of myeloid cells in the brain (both brain-resident microglia and potentially infiltrating monocytes), we quantified the number of Iba1-positive cells in the LC by immunofluorescence. At 14-mo, there was a significant decrease in the number of Iba1-expressing cells in the LC of Tg animals (Fig. 10A,B; t_(13)_=2.845, p=0.0138; n=7), with no changes at other ages. To determine astrocyte activation, we quantified glial fibrillary acidic protein (GFAP) IR, commonly used as a protein marker of astrogliosis (Eng and Ghirnikar, 1994), in the LC. At 24-mo, there was a significant increase in astrocytic GFAP expression in the LC of Tg animals (Fig. 10C, D; t_(10)_=2.744, p=0.0207; n= 7 nTg & 4 Tg).

**Fig 10.**
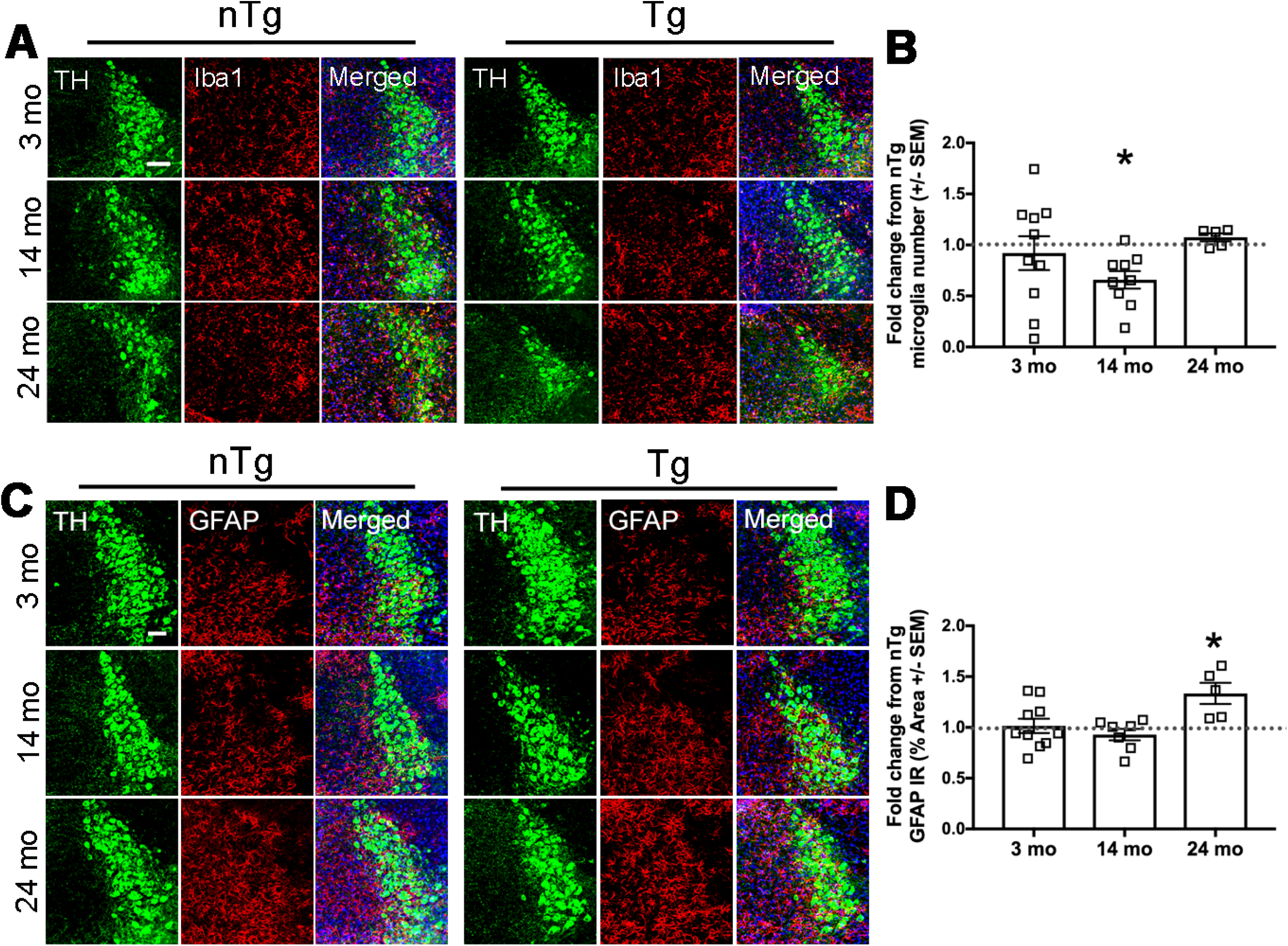
Human asyn in LC neurons affects expression of local inflammatory markers. There are fewer Iba1-positive cells (red) in the LC (TH: green) at 14-mos. ***A,*** Representative immunofluorescent images. ***B***, Microglial count in the LC graphed as fold change Tg from nTg Iba1+ cell count ± SEM. Student’s t-test of nTg and Tg for each age group. Expression of astrocytic GFAP (red) is increased in the Tg LC (TH: green) at 24-mos of age. ***C,*** Representative immunofluorescent images. ***D***, Quantification of GFAP % IR in the LC as percent ROI graphed as fold change Tg IR from nTg mean IR ± SEM. Student’s t-test of nTg and Tg for each age group. Scale bar = 50μm. *\*p*<0.05.

### Hippocampal astrogliosis and changes in number of hippocampal Iba1-expressing cells in DBH-hSNCA mice

To determine whether degeneration of hippocampal LC projections is associated with inflammation, GFAP IR was visualized in the CA1, CA3, and dentate gyrus regions of the hippocampus. At 14-mos, there was a significant increase in GFAP expression in CA1 (Fig. 11A,B; t_(19)_=2.723, p=0.0135, n= 12 nTg & 9 Tg) and CA3 (Fig. 11E,F; t_(19)_=2.275, p=0.0347, n = 12 nTg & 9 Tg) but not the dentate gyrus (Fig. 11C,D; t_(19)_=1.607, p=0.1246, n= 12 nTg & 12 Tg) of Tg mice compared to nTg. Similar to what we observed in the LC, the number of Iba1-expressing cells in CA1 was reduced in Tg mice at 14-mos (Fig. 12; t_(16)_=2.592, p=0.0196, n=9).

**Fig 11:**
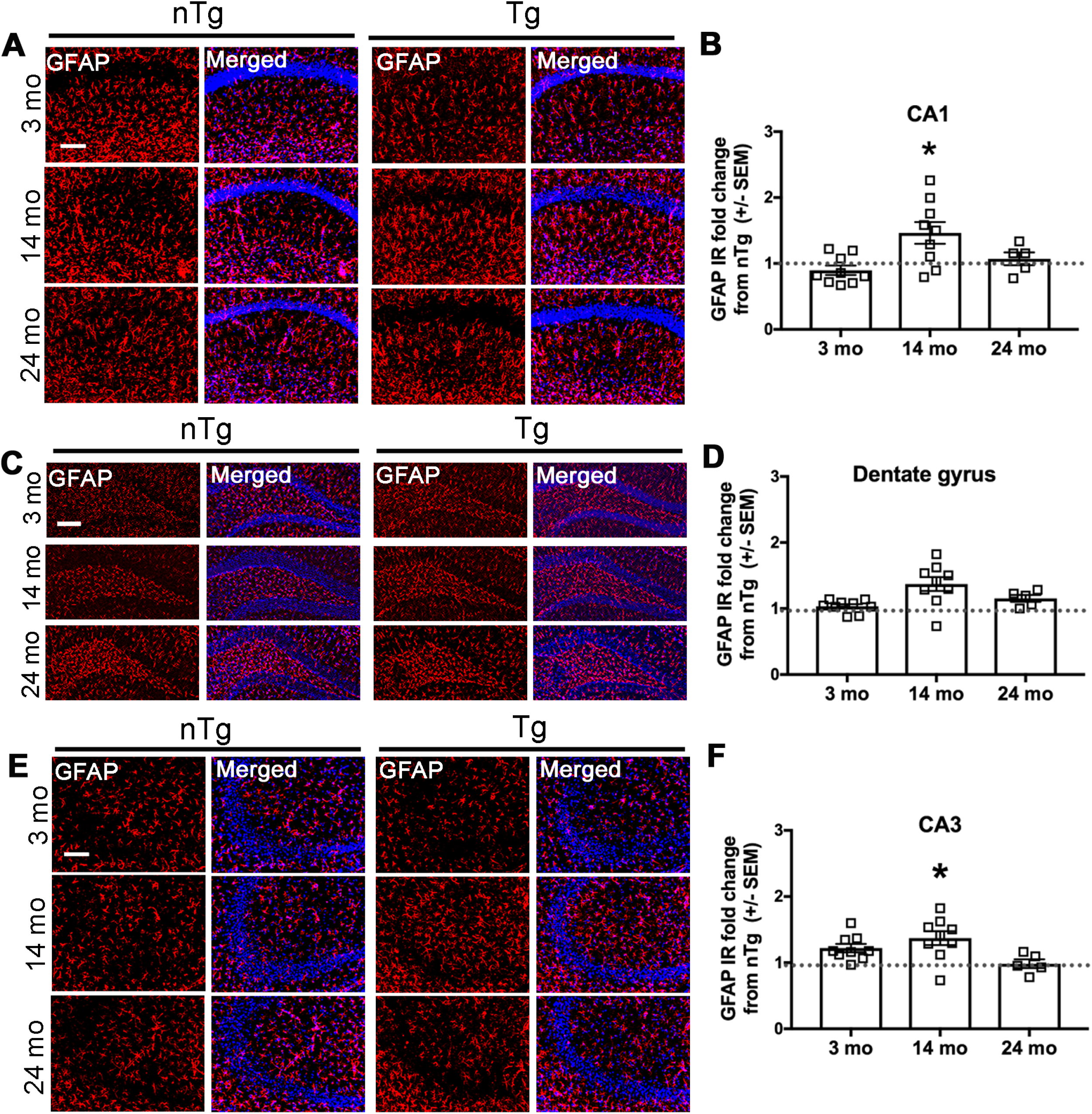
Hippocampal astrocytic GFAP expression is increased at 14-mos. At 14-mos, Tg mice have significantly more GFAP (red) expression than nTg in the CA1 region of the hippocampus. Representative immunofluorescent images of ***A,*** CA1, ***C,*** dentate gyrus, and ***E,*** CA3 regions. Quantification of GFAP IR as % area of ROI in ***B,*** CA1, ***D,*** dentate gyrus, and ***F,*** CA3. Student’s t-test of nTg and Tg for each age group, graphed as fold change Tg from nTg mean ± SEM. Scale bar = 50μm. *\*p*<0.05.

**Fig 12:**
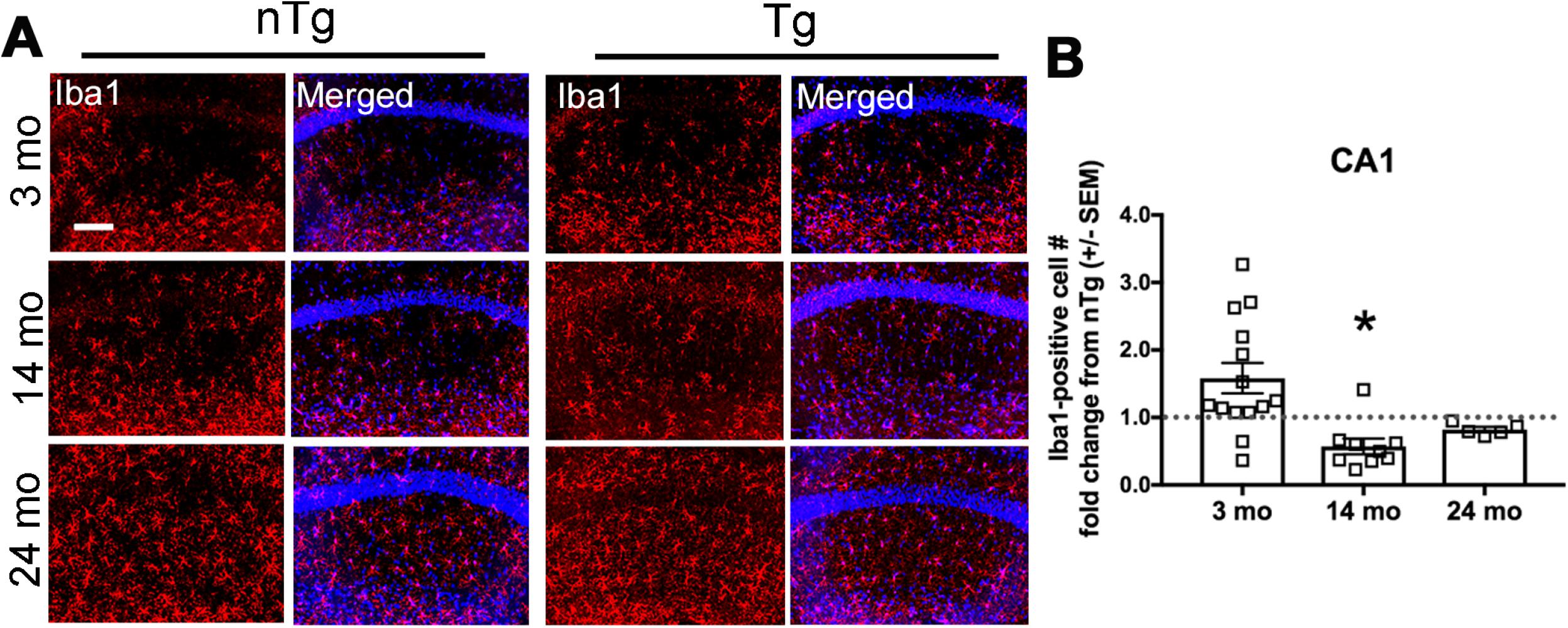
Fewer Iba1-expressing cells in hippocampal CA1 region in 14-mo old Tg DBH-hSNCA mice. At 14-mos, Tg mice have fewer Iba1-expressing cells in hippocampal region CA1. ***A,*** Representative immunofluorescent images of Iba1-expressing cells (red). ***B,*** Quantification of Iba1-expressing cells in CA1 graphed as Tg fold change from nTg mean ± SEM. Student’s t-test by genotype for each age group. Nuclear stain in blue. Scale bar = 150μm. *\*p*<0.05.

### Human asyn expression in LC neurons is associated with loss of hippocampal LC fibers at 24-mos

The LC is the sole source of hippocampal NE, which is necessary for proper memory formation and retrieval (Devauges and Sara, 1991). Noradrenergic LC fibers express NET, and PD brain tissue shows substantial LC denervation (Pavese et al., 2011). To determine whether hippocampal LC projections degenerate in DBH-hSNCA mice, we examined NET IR in the CA1, CA3, and dentate gyrus regions of the hippocampus. At 24-mos, we found a reduction in LC fibers in the dentate gyrus (Fig. 13C,D; t_(10)_=2.974, p=0.0156; n= 6 nTg & 5 Tg), with a trend for reduction in CA1 (Fig. 13A, B; t_(10)_=1.899, p=0.0901, n= 6 nTg & 5 Tg) and CA3 (Fig. 13E,F; t_(10)_=1.538, p=0.1585, n= 6 nTg & 5 Tg). No differences were observed in mice at 3-, or 14-mos.

**Figure 13:**
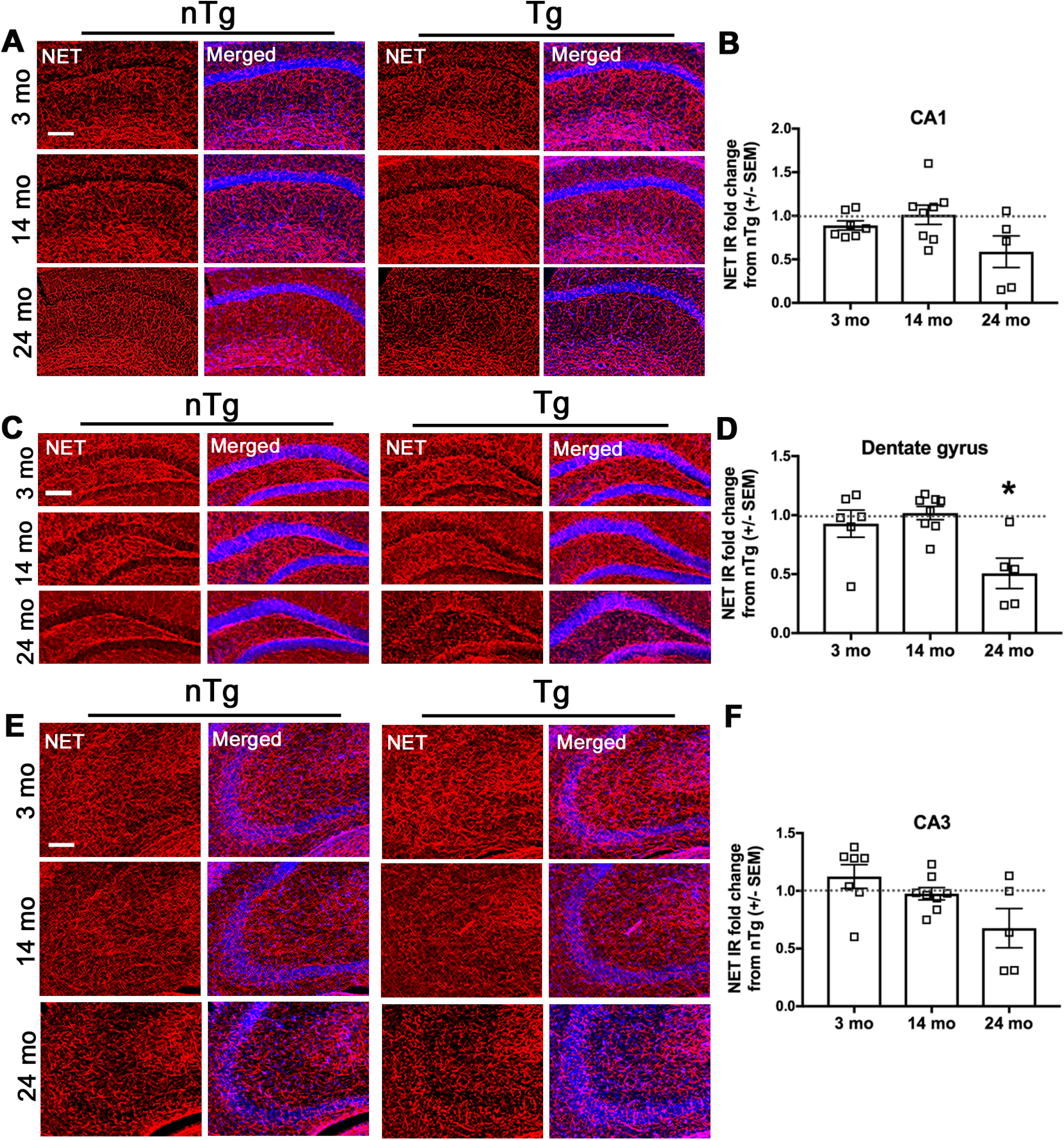
Asyn expression in LC neurons results in age-dependent degeneration of hippocampal LC fibers. NET (red) IR is reduced in the dentate gyrus of Tg mice at 24-mos. Representative immunofluorescent images of ***A,*** CA1, ***C,*** dentate gyrus, and ***E,*** CA3 regions. Quantification of NET IR as percent area of ROI in ***B,*** CA1, ***D,*** dentate gyrus, and ***F,*** CA3. Student’s t-test of nTg and Tg for each age group, graphed as fold change Tg from nTg mean ± SEM. Scale bar = 150μm. *\*p*<0.05.

## Discussion

Based on a wealth of evidence (Mavridis et al., 1991; Srinivasan and Schmidt, 2003; Tong et al., 2006; Rommelfanger et al., 2007; Yao et al., 2015), we posit that asyn pathology and degeneration of LC neurons may represent a tipping point in PD progression; therefore, understanding how asyn accumulation in LC neurons affects their function and survival may help inform development of new therapeutics for earlier interventions. To this end, we developed a new BAC transgenic mouse expressing human wild-type asyn under the control of the noradrenergic-specific DBH.

Human asyn was detectable by immunofluorescence in the Tg LC and in neurons of some (e.g. A4, A5) but not other (e.g. A1, A2) additional DBH-expressing brainstem nuclei by immunofluorescence at 3-mos. Analysis of total asyn (mouse + human) by western blot revealed that the total asyn burden is increased in Tg LC neurons at 3-mos. While pSer129-positive and fibrillar pathological inclusions were not detected in LC neurons at 24-mos, increasing the burden of asyn expression in LC neurons resulted in the formation of putative asyn oligomers, as detected by PLA. We did find nonuniform biochemical and immunohistolocigal evidence of conformation-specific asyn-IR (MJFR14) in 3- and 24-month old Tg mice, suggesting a change in the structural conformation of asyn in some animals, possibly as a consequence of human asyn burden, the corruption of endogenous asyn, or another critical intrinsic, animal-specific factor such as altered TH, which a trend was found to be negatively correlated to the level of conformation-specific asyn by dot blot in 3-month old Tg mice. Although we observed no loss of LC cell bodies, NET-expressing fibers were reduced in the hippocampus of 24-mo old mice. This finding resembles the pattern of neuron death observed in PD, with axon terminals degenerating prior to frank cell loss in the LC (Hornykiewicz, 1998). Interestingly, the selective loss of LC fibers in the dentate gyrus of DBH-hSNCA mice is reminiscent of what we observed in TgF344-AD rat model of Alzheimer’s disease that accumulates tau pathology in the LC (Rorabaugh et al., 2017). To determine the functional outcome of human asyn expression in LC neurons, we measured catecholamine levels in the hippocampus and striatum. Despite the loss of hippocampal LC fibers at 24-mos, we did not detect changes in hippocampal NE content by HPLC, which may indicate a compensatory enhancement in NE synthesis. We observed a trend toward decreased striatal NE 24-mos, but it was not statistically significant. With the sparse noradrenergic innervation to the striatum, it is possible that changes in NE neurotransmission are detectable there prior to more densely innervated regions. Dysregulated DA metabolism is a central feature of PD (Leenders et al., 1990), and midbrain dopaminergic innervation to the striatum is modulated by LC-NE (Lategan et al., 1990; Grenhoff et al., 1993; Rommelfanger et al., 2007; Rommelfanger and Weinshenker, 2007). Studies of catecholamine function in rodents show that enhancing LC-NE stimulates midbrain DA release in the striatum, whereas LC lesions or NE deficiency reduce striatal DA release (Lategan et al., 1990; Grenhoff et al., 1993; Schank et al., 2006). We found that at 24-mos, Tg mice had a reduced striatal ratio of DOPAC:DA and a trend toward increased DA, suggesting that human asyn expression in aging LC neurons causes a reduction in striatal DA turnover. Future studies will determine whether NE innervation to midbrain DA neurons is reduced in 24-mo DBH-hSNCA mice, and whether loss of noradrenergic transmission impairs striatal DA release. While no significant differences were detected in hippocampal or striatal NE content at any age, Tg LC neurons had increased TH expression at 3- and 14-mos relative to nTg littermates, suggesting an increased capacity for NE synthesis.

Neuroinflammation is a central feature of PD pathology (McGeer et al., 1988; Gerhard et al., 2006; Tansey and Goldberg, 2010), with extensive evidence of changes in microglial activation in brain regions that degenerate in PD (Kim and Joh, 2006; Tansey and Goldberg, 2010), and we know that microglia and astrocyte functions are modulated by NE neurotransmission (Fahrig, 1993; Heneka et al., 2010; Bharani et al., 2017). Unexpectedly, we observed fewer Iba1-expressing cells in the LC and CA1 region of the hippocampus of DBH-hSNCA mice at 14-mos compared to nTg littermates. It has been reported that NE can modulate microglial activity, and the functional outcome of microglia adrenergic receptor activation appears to depend on the physiological context (Fujita et al., 1998; Gyoneva and Traynelis, 2013; Johnson et al., 2013). At 14-mos, there was an increase in hippocampal astrocytic GFAP expression. Astrocytes throughout the brain are activated by NE, and AR signaling stimulates calcium transients and promotes the release of inflammatory signaling molecules by astrocytes (Norris and Benveniste, 1993; Duffy and MacVicar, 1995; Paukert et al., 2014). However, the number of GFAP-expressing astrocytes is inversely correlated to the amount of dopaminergic cell loss in PD, and increased GFAP-IR in the hippocampus occurred prior to loss of LC projections (Damier et al., 1993; Eng and Ghirnikar, 1994). Further studies are required to determine whether increased GFAP activity is due to dysregulated NE neurotransmission, or an inflammatory response secondary to degeneration of LC projections.

Historically, the LC has been implicated in arousal state and stress responses. For example, LC activity tracks with sleep cycles (with highest firing during wake and immediately preceding sleep-wake transitions), and chemogenetic or optogenetic activation of LC neurons increases wakefulness (Carter et al., 2010; Vazey and Aston-Jones, 2014; Porter-Stransky et al., 2019). Furthermore, the LC is activated by stress, and stimulation of LC neurons elicits anxiety-like behaviors (Valentino and Van Bockstaele, 2008; McCall et al., 2015). DBH-hSNCA mice exhibited age-dependent behavioral phenotypes that peaked at 14-mos and are consistent with LC hyperactivity and increased NE transmission. Specifically, compared to nTg littermates, DBH-hSNCA mice displayed increased arousal (as measured by latency to fall asleep), anxiety-like behavior (as measured by marble burying and latency to re-enter the center of an open field), and stress responses (as measured by freezing during fear conditioning training and context re-exposure). These phenotypes are relevant to multiple non-motor symptoms of PD. Sleep disturbances are one of the most common complaints of PD patients, and patients who experience disturbed sleep appear to have greater LC asyn pathology than those who do not report sleep disturbances (Kalaitzakis et al., 2013). Anxiety is also a common complaint, as up to 60% of PD patients report experiencing anxiety (Chaudhuri and Schapira, 2009; Lin et al., 2015). Importantly, DBH-hSNCA mice did not display changes in the number or speed of ambulations relative to nTg mice at any age examined, ruling out a general locomotor abnormality that has been observed in more ubiquitous asyn overexpression mice (Giasson et al., 2002; Fleming et al., 2004; Graham and Sidhu, 2010). Reducing DA specifically in DA-neurons causes motor, but not most non-motor, deficits in mice, which further indicates that these symptoms arise from alterations in distinct neurotransmitter systems (Jiang et al., 2020). In fact, the increased sleep latency observed in Tg animals was normalized by blocking α1- and β-adrenergic receptors, which suggests that the behavioral phenotype is due to enhanced noradrenergic neurotransmission. We have previously speculated that asyn pathology promotes LC hyperactivity and non-motor symptoms during PD progression prior to the degeneration of noradrenergic neurons later in the disease (Weinshenker, 2018), and the present data support that idea. Indeed, the behavioral phenotypes we observed were most prominent at 14-mos, while degeneration of LC fibers was not evident until 24-mos, a time when behavioral abnormalities abated. There are several potential mechanisms by which increases in human asyn may be affecting the LC-NE system (Fig. 14). TH protein was increased in 3- and 14-mo old Tg mice, which is when behavioral phenotypes consistent with NE over-activity emerged. Because TH is the rate-limiting enzyme in NE production, this may reflect increased NE synthetic capacity, although the relevance is not clear because we did not detect differences in tissue NE content. Human wild-type asyn has been reported to increase in the size, and delay in the closing, of the vesicular fusion pore, allowing more neurotransmitter to spill into the extracellular space (Larsen et al., 2006; Logan et al., 2017). In addition, viral-mediated overexpression of A53T mutant asyn in LC neurons causes increased firing rate (Henrich et al., 2018). Alternatively, if human asyn expression is negatively impacting LC neuron health as suggested by the presence of asyn oligomers and the loss of hippocampal LC fibers at 24-mos, there may be compensatory increases in neuronal activity. Chemical lesion of LC neurons transiently increases NE neurotransmission by increasing neuronal firing frequency and activity patterns (Szot et al., 2016). Future studies directly examining LC neuron firing and NE release will be necessary to identify the mechanism in play here.

**Figure 14:**
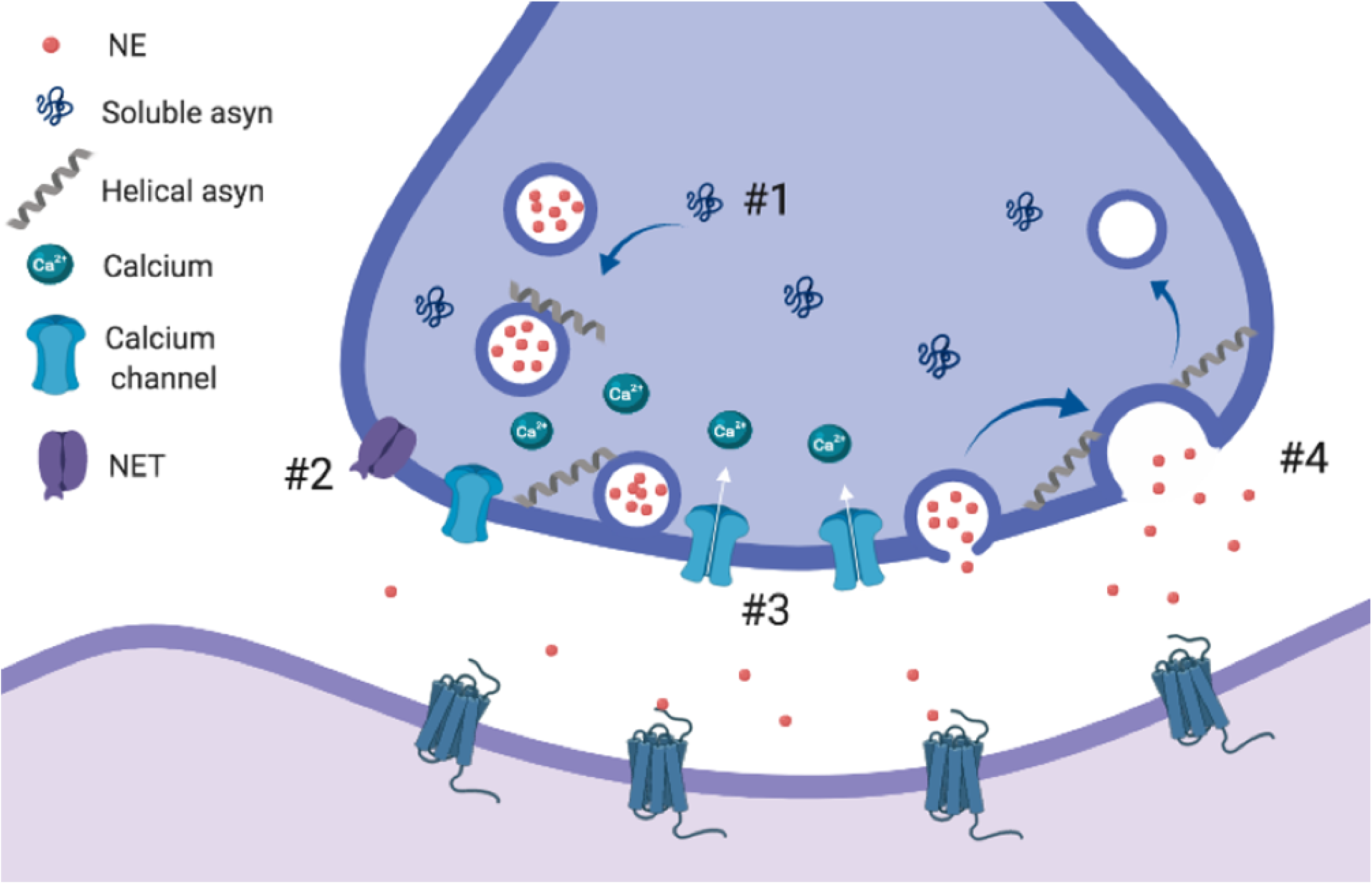
Proposed model of human asyn effects on noradrenergic neurotransmission. Human asyn may enhance extracellular NE by several mechanisms. #1 – Increased expression of asyn could potentiate clustering of synaptic vesicles at the presynaptic membrane. #2 – Elevated asyn expression could reduce norepinephrine transporter (NET) expression at the plasma membrane. #3 – asyn modulation of L-type voltage gated calcium channels could enhance neuronal excitability. #4 – human asyn expression could enlarge the size of and slow the closing of vesicular fusion pores, allowing more NE to spill into the extracellular space.

LC neurons are among the first affected in PD, and the features of DBH-hSNCA mice may represent early pathology and non-motor components of PD that provide insight into the functional impact of human asyn expression in LC neurons during aging. Importantly, our data suggest that overexpression of wild-type human asyn and formation of oligomers alone is sufficient for the LC dysfunction, fiber degeneration, and behavioral deficits we observed, while asyn pSer129 inclusions are not required for these phenotypes. Additionally, studies involving exposure to environmental factors that synergize with asyn expression to influence the risk of PD will likely elucidate the genetic and environmental interactions that contribute to LC involvement in the pre-clinical pre-motor stages of PD, as well as the impact of LC degeneration on the trajectory of PD pathogenesis.

## Acknowledgements

We thank Terina Martinez, Tim Sampson, and the Tansey, Weinshenker, and Giasson labs for useful discussions. This work was supported by NIH/NINDS 5F31 NS098673 (LB), NIH/NINDS F32 NS098615 (KP), NIH/NINDS 1R01 NS102306 (DW), NIH/NIA 1RF1 AG047667 (DW), NIH/NIA 1R01AG061175 (DW), NIH/NINDS 1R01 NS100876 (BG), NIH/NIA 1R01 AG057247 (MGT), NIH/NINDS 5R01 NS092122 (MGT), NIH/NIA 3RF1 AG051514-01 (MGT), and the Norman Fixel Institute for Neurological Diseases (MGT). This research project was also supported in part by the Emory University Integrated Cellular Imaging Microscopy Core, by the Rodent Behavioral Core (RBC), which is subsidized by the Emory University School of Medicine and is one of the Emory Integrated Core Facilities. Additional support was provided by the Emory Neuroscience NINDS Core Facilities P30NS055077. Further support was provided by the Georgia Clinical & Translational Science Alliance of the National Institutes of Health under Award Number UL1TR002378. This study was supported in part by the Emory HPLC Bioanalytical Core (EHBC), which was supported by the Department of Pharmacology, Emory University School of Medicine and the Georgia Clinical & T ranslational Science Alliance of the National Institutes of Health under Award Number UL1TR002378. The content is solely the responsibility of the authors and does not necessarily reflect the official views of the National Institutes of Health.

## Notes

### Competing Interest Statement

The authors have declared no competing interest.

### Summary of Updates

New figures which now include pharmacological data supporting the role of noradrenergic transmission in the behavioral phenotypes (Figure 5), the presence of oligomerc and conformation-specific alpha-synuclein using three independent assays in the absence of mature phosphorylated inclusions (Figure 6), and a proposed model by which increased human asyn may interfere with noradrenergic neurotransmission (Figure 14).

## References

Abbott RD, Ross GW, White LR, Tanner CM, Masaki KH, Nelson JS, Curb JD, Petrovitch H (2005) Excessive daytime sleepiness and subsequent development of Parkinson disease. Neurology 65:1442–1446.

Baekelandt V, Claeys A, Eggermont K, Lauwers E, De Strooper B, Nuttin B, Debyser Z (2002) Characterization of lentiviral vector-mediated gene transfer in adult mouse brain. Hum Gene Ther 13:841–853.

Bharani KL, Derex R, Granholm AC, Ledreux A (2017) A noradrenergic lesion aggravates the effects of systemic inflammation on the hippocampus of aged rats. PLoS One 12:e0189821.

Braak E, Sandmann-Keil D, Rub U, Gai WP, de Vos RA, Steur EN, Arai K, Braak H (2001) alpha-synuclein immunopositive Parkinson’s disease-related inclusion bodies in lower brain stem nuclei. Acta Neuropathol 101:195–201.

Britton DR, Britton KT (1981) A sensitive open field measure of anxiolytic drug activity. Pharmacol Biochem Behav 15:577–582.

Brunnstrom H, Friberg N, Lindberg E, Englund E (2011) Differential degeneration of the locus coeruleus in dementia subtypes. Clin Neuropathol 30:104–110.

Butkovich LM, Houser MC, Tansey MG (2018) alpha-Synuclein and Noradrenergic Modulation of Immune Cells in Parkinson’s Disease Pathogenesis. Front Neurosci 12:626.

Carter ME, Yizhar O, Chikahisa S, Nguyen H, Adamantidis A, Nishino S, Deisseroth K, de Lecea L (2010) Tuning arousal with optogenetic modulation of locus coeruleus neurons. Nat Neurosci 13:1526–1533.

Chalermpalanupap T, Schroeder JP, Rorabaugh JM, Liles LC, Lah JJ, Levey AI, Weinshenker D (2018) Locus Coeruleus Ablation Exacerbates Cognitive Deficits, Neuropathology, and Lethality in P301S Tau Transgenic Mice. J Neurosci 38:74–92.

Chartier-Harlin MC, Kachergus J, Roumier C, Mouroux V, Douay X, Lincoln S, Levecque C, Larvor L, Andrieux J, Hulihan M, Waucquier N, Defebvre L, Amouyel P, Farrer M, Destee A (2004) Alpha-synuclein locus duplication as a cause of familial Parkinson’s disease. Lancet 364:1167–1169.

Chaudhuri KR, Schapira AH (2009) Non-motor symptoms of Parkinson’s disease: dopaminergic pathophysiology and treatment. Lancet Neurol 8:464–474.

Chen X, Huddleston DE, Langley J, Ahn S, Barnum CJ, Factor SA, Levey AI, Hu X (2014) Simultaneous imaging of locus coeruleus and substantia nigra with a quantitative neuromelanin MRI approach. Magn Reson Imaging 32:1301–1306.

Chui HC, Mortimer JA, Slager U, Zarow C, Bondareff W, Webster DD (1986) Pathologic correlates of dementia in Parkinson’s disease. Arch Neurol 43:991–995.

Cubells JF, Schroeder JP, Barrie ES, Manvich DF, Sadee W, Berg T, Mercer K, Stowe TA, Liles LC, Squires KE, Mezher A, Curtin P, Perdomo DL, Szot P, Weinshenker D (2016) Human Bacterial Artificial Chromosome (BAC) Transgenesis Fully Rescues Noradrenergic Function in Dopamine beta-Hydroxylase Knockout Mice. PLoS One 11:e0154864.

Damier P, Hirsch EC, Zhang P, Agid Y, Javoy-Agid F (1993) Glutathione peroxidase, glial cells and Parkinson’s disease. Neuroscience 52:1–6.

de Sousa Rodrigues ME, Bekhbat M, Houser MC, Chang J, Walker DI, Jones DP, Oller do Nascimento CMP, Barnum CJ, Tansey MG (2017) Chronic psychological stress and high-fat high-fructose diet disrupt metabolic and inflammatory gene networks in the brain, liver, and gut and promote behavioral deficits in mice. Brain Behav Immun 59:158–172.

Delenclos M, Faroqi AH, Yue M, Kurti A, Castanedes-Casey M, Rousseau L, Phillips V, Dickson DW, Fryer JD, McLean PJ (2017) Neonatal AAV delivery of alpha-synuclein induces pathology in the adult mouse brain. Acta Neuropathol Commun 5:51.

den Hartog JW, Bethlem J (1960) The distribution of Lewy bodies in the central and autonomic nervous systems in idiopathic paralysis agitans. J Neurol Neurosurg Psychiatry 23:283–290.

Devauges V, Sara SJ (1991) Memory retrieval enhancement by locus coeruleus stimulation: evidence for mediation by beta-receptors. Behav Brain Res 43:93–97.

Duffy S, MacVicar BA (1995) Adrenergic calcium signaling in astrocyte networks within the hippocampal slice. J Neurosci 15:5535–5550.

Eng LF, Ghirnikar RS (1994) GFAP and astrogliosis. Brain Pathol 4:229–237.

Fahrig T (1993) Receptor subtype involved and mechanism of norepinephrine-induced stimulation of glutamate uptake into primary cultures of rat brain astrocytes. Glia 7:212–218.

Fearnley JM, Lees AJ (1991) Ageing and Parkinson’s disease: substantia nigra regional selectivity. Brain 114 (Pt 5):2283–2301.

Ferese R, Modugno N, Campopiano R, Santilli M, Zampatti S, Giardina E, Nardone A, Postorivo D, Fornai F, Novelli G, Romoli E, Ruggieri S, Gambardella S (2015) Four Copies of SNCA Responsible for Autosomal Dominant Parkinson’s Disease in Two Italian Siblings. Parkinsons Dis 2015:546462.

Fleming SM, Salcedo J, Fernagut PO, Rockenstein E, Masliah E, Levine MS, Chesselet MF (2004) Early and progressive sensorimotor anomalies in mice overexpressing wild-type human alpha-synuclein. J Neurosci 24:9434–9440.

Fujita H, Tanaka J, Maeda N, Sakanaka M (1998) Adrenergic agonists suppress the proliferation of microglia through beta 2-adrenergic receptor. Neurosci Lett 242:37–40.

Fujiwara H, Hasegawa M, Dohmae N, Kawashima A, Masliah E, Goldberg MS, Shen J, Takio K, Iwatsubo T (2002) alpha-Synuclein is phosphorylated in synucleinopathy lesions. Nat Cell Biol 4:160–164.

Gerhard A, Pavese N, Hotton G, Turkheimer F, Es M, Hammers A, Eggert K, Oertel W, Banati RB, Brooks DJ (2006) In vivo imaging of microglial activation with [11C](R)-PK11195 PET in idiopathic Parkinson’s disease. Neurobiol Dis 21:404–412.

German DC, Manaye KF, White CL, 3rd, Woodward DJ, McIntire DD, Smith WK, Kalaria RN, Mann DM (1992) Disease-specific patterns of locus coeruleus cell loss. Ann Neurol 32:667–676.

Giasson BI, Duda JE, Quinn SM, Zhang B, Trojanowski JQ, Lee VM (2002) Neuronal alpha-synucleinopathy with severe movement disorder in mice expressing A53T human alpha-synuclein. Neuron 34:521–533.

Gonera EG, van’t Hof M, Berger HJ, van Weel C, Horstink MW (1997) Symptoms and duration of the prodromal phase in Parkinson’s disease. Mov Disord 12:871–876.

Graham DR, Sidhu A (2010) Mice expressing the A53T mutant form of human alpha-synuclein exhibit hyperactivity and reduced anxiety-like behavior. J Neurosci Res 88:1777–1783.

Grenhoff J, Nisell M, Ferre S, Aston-Jones G, Svensson TH (1993) Noradrenergic modulation of midbrain dopamine cell firing elicited by stimulation of the locus coeruleus in the rat. J Neural Transm Gen Sect 93:11–25.

Gyoneva S, Traynelis SF (2013) Norepinephrine modulates the motility of resting and activated microglia via different adrenergic receptors. J Biol Chem 288:15291–15302.

Halliday GM, Li YW, Blumbergs PC, Joh TH, Cotton RG, Howe PR, Blessing WW, Geffen LB (1990) Neuropathology of immunohistochemically identified brainstem neurons in Parkinson’s disease. Ann Neurol 27:373–385.

Hansen C, Bjorklund T, Petit GH, Lundblad M, Murmu RP, Brundin P, Li JY (2013) A novel alpha-synuclein-GFP mouse model displays progressive motor impairment, olfactory dysfunction and accumulation of alpha-synuclein-GFP. Neurobiol Dis 56:145–155.

Haycock JW (1993) Multiple signaling pathways in bovine chromaffin cells regulate tyrosine hydroxylase phosphorylation at Ser19, Ser31, and Ser40. Neurochem Res 18:15–26.

Heneka MT, Nadrigny F, Regen T, Martinez-Hernandez A, Dumitrescu-Ozimek L, Terwel D, Jardanhazi-Kurutz D, Walter J, Kirchhoff F, Hanisch UK, Kummer MP (2010) Locus ceruleus controls Alzheimer’s disease pathology by modulating microglial functions through norepinephrine. Proc Natl Acad Sci U S A 107:6058–6063.

Henrich MT, Geibl FF, Lee B, Chiu WH, Koprich JB, Brotchie JM, Timmermann L, Decher N, Matschke LA, Oertel WH (2018) A53T-alpha-synuclein overexpression in murine locus coeruleus induces Parkinson’s disease-like pathology in neurons and glia. Acta Neuropathol Commun 6:39.

Hirsch E, Graybiel AM, Agid YA (1988) Melanized dopaminergic neurons are differentially susceptible to degeneration in Parkinson’s disease. Nature 334:345–348.

Hobson JA, McCarley RW, Wyzinski PW (1975) Sleep cycle oscillation: reciprocal discharge by two brainstem neuronal groups. Science 189:55–58.

Hornykiewicz O (1998) Biochemical aspects of Parkinson’s disease. Neurology 51:S2–9.

Hunsley MS, Palmiter RD (2004) Altered sleep latency and arousal regulation in mice lacking norepinephrine. Pharmacol Biochem Behav 78:765–773.

Ip CW, Klaus LC, Karikari AA, Visanji NP, Brotchie JM, Lang AE, Volkmann J, Koprich JB (2017) AAV1/2-induced overexpression of A53T-alpha-synuclein in the substantia nigra results in degeneration of the nigrostriatal system with Lewy-like pathology and motor impairment: a new mouse model for Parkinson’s disease. Acta Neuropathol Commun 5:11.

Iversen LL, Rossor MN, Reynolds GP, Hills R, Roth M, Mountjoy CQ, Foote SL, Morrison JH, Bloom FE (1983) Loss of pigmented dopamine-beta-hydroxylase positive cells from locus coeruleus in senile dementia of Alzheimer’s type. Neurosci Lett 39:95–100.

Jiang S, Berger S, Hu Y, Bartsch D, Tian Y (2020) Alterations of the Motor and Olfactory Functions Related to Parkinson’s Disease in Transgenic Mice With a VMAT2-Deficiency in Dopaminergic Neurons. Front Neurosci 14:356.

Joers V, Dilley K, Rahman S, Jones C, Shultz J, Simmons H, Emborg ME (2014) Cardiac sympathetic denervation in 6-OHDA-treated nonhuman primates. PLoS One 9:e104850.

Johnson JD, Zimomra ZR, Stewart LT (2013) Beta-adrenergic receptor activation primes microglia cytokine production. J Neuroimmunol 254:161–164.

Kalaitzakis ME, Gentleman SM, Pearce RK (2013) Disturbed sleep in Parkinson’s disease: anatomical and pathological correlates. Neuropathol Appl Neurobiol 39:644–653.

Keren NI, Taheri S, Vazey EM, Morgan PS, Granholm AC, Aston-Jones GS, Eckert MA (2015) Histologic validation of locus coeruleus MRI contrast in post-mortem tissue. Neuroimage 113:235–245.

Kilbourn MR, Sherman P, Abbott LC (1998) Reduced MPTP neurotoxicity in striatum of the mutant mouse tottering. Synapse 30:205–210.

Kim YS, Joh TH (2006) Microglia, major player in the brain inflammation: their roles in the pathogenesis of Parkinson’s disease. Exp Mol Med 38:333–347.

Kirik D, Rosenblad C, Burger C, Lundberg C, Johansen TE, Muzyczka N, Mandel RJ, Bjorklund A (2002) Parkinson-like neurodegeneration induced by targeted overexpression of alpha-synuclein in the nigrostriatal system. J Neurosci 22:2780–2791.

Koprich JB, Johnston TH, Reyes MG, Sun X, Brotchie JM (2010) Expression of human A53T alpha-synuclein in the rat substantia nigra using a novel AAV1/2 vector produces a rapidly evolving pathology with protein aggregation, dystrophic neurite architecture and nigrostriatal degeneration with potential to model the pathology of Parkinson’s disease. Mol Neurodegener 5:43.

Kreiner G, Rafa-Zablocka K, Barut J, Chmielarz P, Kot M, Baginska M, Parlato R, Daniel WA, Nalepa I (2019) Stimulation of noradrenergic transmission by reboxetine is beneficial for a mouse model of progressive parkinsonism. Sci Rep 9:5262.

Kumer SC, Vrana KE (1996) Intricate regulation of tyrosine hydroxylase activity and gene expression. J Neurochem 67:443–462.

Larsen KE, Schmitz Y, Troyer MD, Mosharov E, Dietrich P, Quazi AZ, Savalle M, Nemani V, Chaudhry FA, Edwards RH, Stefanis L, Sulzer D (2006) Alpha-synuclein overexpression in PC12 and chromaffin cells impairs catecholamine release by interfering with a late step in exocytosis. J Neurosci 26:11915–11922.

Lategan AJ, Marien MR, Colpaert FC (1990) Effects of locus coeruleus lesions on the release of endogenous dopamine in the rat nucleus accumbens and caudate nucleus as determined by intracerebral microdialysis. Brain Res 523:134–138.

Leenders KL, Salmon EP, Tyrrell P, Perani D, Brooks DJ, Sager H, Jones T, Marsden CD, Frackowiak RS (1990) The nigrostriatal dopaminergic system assessed in vivo by positron emission tomography in healthy volunteer subjects and patients with Parkinson’s disease. Arch Neurol 47:1290–1298.

Levitt M, Spector S, Sjoerdsma A, Udenfriend S (1965) Elucidation of the Rate-Limiting Step in Norepinephrine Biosynthesis in the Perfused Guinea-Pig Heart. J Pharmacol Exp Ther 148:1–8.

Lin CH, Lin JW, Liu YC, Chang CH, Wu RM (2015) Risk of Parkinson’s disease following anxiety disorders: a nationwide population-based cohort study. Eur J Neurol 22:1280–1287.

Lloyd GM, Trejo-Lopez JA, Xia Y, McFarland KN, Lincoln SJ, Ertekin-Taner N, Giasson BI, Yachnis AT, Prokop S (2020) Prominent amyloid plaque pathology and cerebral amyloid angiopathy in APP V717I (London) carrier - phenotypic variability in autosomal dominant Alzheimer’s disease. Acta Neuropathol Commun 8:31.

Logan T, Bendor J, Toupin C, Thorn K, Edwards RH (2017) alpha-Synuclein promotes dilation of the exocytotic fusion pore. Nat Neurosci 20:681–689.

Mann JJ, Stanley M, Kaplan RD, Sweeney J, Neophytides A (1983) Central catecholamine metabolism in vivo and the cognitive and motor deficits in Parkinson’s disease. J Neurol Neurosurg Psychiatry 46:905–910.

Maskri L, Zhu X, Fritzen S, Kuhn K, Ullmer C, Engels P, Andriske M, Stichel CC, Lubbert H (2004) Influence of different promoters on the expression pattern of mutated human alpha-synuclein in transgenic mice. Neurodegener Dis 1:255–265.

Masliah E, Rockenstein E, Veinbergs I, Mallory M, Hashimoto M, Takeda A, Sagara Y, Sisk A, Mucke L (2000) Dopaminergic loss and inclusion body formation in alpha-synuclein mice: implications for neurodegenerative disorders. Science 287:1265–1269.

Mavridis M, Degryse AD, Lategan AJ, Marien MR, Colpaert FC (1991) Effects of locus coeruleus lesions on parkinsonian signs, striatal dopamine and substantia nigra cell loss after 1-methyl-4-phenyl-1,2,3,6-tetrahydropyridine in monkeys: a possible role for the locus coeruleus in the progression of Parkinson’s disease. Neuroscience 41:507–523.

McCall JG, Al-Hasani R, Siuda ER, Hong DY, Norris AJ, Ford CP, Bruchas MR (2015) CRH Engagement of the Locus Coeruleus Noradrenergic System Mediates Stress-Induced Anxiety. Neuron 87:605–620.

McGeer PL, Itagaki S, Boyes BE, McGeer EG (1988) Reactive microglia are positive for HLA-DR in the substantia nigra of Parkinson’s and Alzheimer’s disease brains. Neurology 38:1285–1291.

Murchison CF, Zhang XY, Zhang WP, Ouyang M, Lee A, Thomas SA (2004) A distinct role for norepinephrine in memory retrieval. Cell 117:131–143.

Niu H, Shen L, Li T, Ren C, Ding S, Wang L, Zhang Z, Liu X, Zhang Q, Geng D, Wu X, Li H (2018) Alpha-synuclein overexpression in the olfactory bulb initiates prodromal symptoms and pathology of Parkinson’s disease. Transl Neurodegener 7:25.

Norris JG, Benveniste EN (1993) Interleukin-6 production by astrocytes: induction by the neurotransmitter norepinephrine. J Neuroimmunol 45:137–145.

Paukert M, Agarwal A, Cha J, Doze VA, Kang JU, Bergles DE (2014) Norepinephrine controls astroglial responsiveness to local circuit activity. Neuron 82:1263–1270.

Pavese N, Rivero-Bosch M, Lewis SJ, Whone AL, Brooks DJ (2011) Progression of monoaminergic dysfunction in Parkinson’s disease: a longitudinal 18F-dopa PET study. Neuroimage 56:1463–1468.

Phillips RG, LeDoux JE (1992) Differential contribution of amygdala and hippocampus to cued and contextual fear conditioning. Behav Neurosci 106:274–285.

Pifl C, Kish SJ, Hornykiewicz O (2012) Thalamic noradrenaline in Parkinson’s disease: deficits suggest role in motor and non-motor symptoms. Mov Disord 27:1618–1624.

Porter-Stransky KA, Centanni SW, Karne SL, Odil LM, Fekir S, Wong JC, Jerome C, Mitchell HA, Escayg A, Pedersen NP, Winder DG, Mitrano DA, Weinshenker D (2019) Noradrenergic Transmission at Alpha1-Adrenergic Receptors in the Ventral Periaqueductal Gray Modulates Arousal. Biol Psychiatry 85:237–247.

Remy P, Doder M, Lees A, Turjanski N, Brooks D (2005) Depression in Parkinson’s disease: loss of dopamine and noradrenaline innervation in the limbic system. Brain 128:1314–1322.

Rinaman L (2011) Hindbrain noradrenergic A2 neurons: diverse roles in autonomic, endocrine, cognitive, and behavioral functions. Am J Physiol Regul Integr Comp Physiol 300:R222–235.

Rommelfanger KS, Weinshenker D (2007) Norepinephrine: The redheaded stepchild of Parkinson’s disease. Biochem Pharmacol 74:177–190.

Rommelfanger KS, Weinshenker D, Miller GW (2004) Reduced MPTP toxicity in noradrenaline transporter knockout mice. J Neurochem 91:1116–1124.

Rommelfanger KS, Edwards GL, Freeman KG, Liles LC, Miller GW, Weinshenker D (2007) Norepinephrine loss produces more profound motor deficits than MPTP treatment in mice. Proc Natl Acad Sci U S A 104:13804–13809.

Rorabaugh JM, Chalermpalanupap T, Botz-Zapp CA, Fu VM, Lembeck NA, Cohen RM, Weinshenker D (2017) Chemogenetic locus coeruleus activation restores reversal learning in a rat model of Alzheimer’s disease. Brain 140:3023–3038.

Ross GW, Petrovitch H, Abbott RD, Tanner CM, Popper J, Masaki K, Launer L, White LR (2008) Association of olfactory dysfunction with risk for future Parkinson’s disease. Ann Neurol 63:167–173.

Rutherford NJ, Brooks M, Giasson BI (2016) Novel antibodies to phosphorylated alpha-synuclein serine 129 and NFL serine 473 demonstrate the close molecular homology of these epitopes. Acta Neuropathol Commun 4:80.

Sampson TR, Debelius JW, Thron T, Janssen S, Shastri GG, Ilhan ZE, Challis C, Schretter CE, Rocha S, Gradinaru V, Chesselet MF, Keshavarzian A, Shannon KM, Krajmalnik-Brown R, Wittung-Stafshede P, Knight R, Mazmanian SK (2016) Gut Microbiota Regulate Motor Deficits and Neuroinflammation in a Model of Parkinson’s Disease. Cell 167:1469–1480 e1412.

Sawamoto K, Nakao N, Kobayashi K, Matsushita N, Takahashi H, Kakishita K, Yamamoto A, Yoshizaki T, Terashima T, Murakami F, Itakura T, Okano H (2001) Visualization, direct isolation, and transplantation of midbrain dopaminergic neurons. Proc Natl Acad Sci U S A 98:6423–6428.

Schank JR, Ventura R, Puglisi-Allegra S, Alcaro A, Cole CD, Liles LC, Seeman P, Weinshenker D (2006) Dopamine beta-hydroxylase knockout mice have alterations in dopamine signaling and are hypersensitive to cocaine. Neuropsychopharmacology 31:2221–2230.

Schell H, Hasegawa T, Neumann M, Kahle PJ (2009) Nuclear and neuritic distribution of serine-129 phosphorylated alpha-synuclein in transgenic mice. Neuroscience 160:796–804.

Singleton AB et al. (2003) alpha-Synuclein locus triplication causes Parkinson’s disease. Science 302:841.

Song CH, Fan X, Exeter CJ, Hess EJ, Jinnah HA (2012) Functional analysis of dopaminergic systems in a DYT1 knock-in mouse model of dystonia. Neurobiol Dis 48:66–78.

Song S, Jiang L, Oyarzabal EA, Wilson B, Li Z, Shih YI, Wang Q, Hong JS (2018) Loss of Brain Norepinephrine Elicits Neuroinflammation-Mediated Oxidative Injury and Selective Caudo-Rostral Neurodegeneration. Mol Neurobiol.

Sotiriou E, Vassilatis DK, Vila M, Stefanis L (2010) Selective noradrenergic vulnerability in alpha-synuclein transgenic mice. Neurobiol Aging 31:2103–2114.

Spillantini MG, Schmidt ML, Lee VM, Trojanowski JQ, Jakes R, Goedert M (1997) Alpha-synuclein in Lewy bodies. Nature 388:839–840.

Srinivasan J, Schmidt WJ (2003) Potentiation of parkinsonian symptoms by depletion of locus coeruleus noradrenaline in 6-hydroxydopamine-induced partial degeneration of substantia nigra in rats. Eur J Neurosci 17:2586–2592.

Szot P, Franklin A, Miguelez C, Wang Y, Vidaurrazaga I, Ugedo L, Sikkema C, Wilkinson CW, Raskind MA (2016) Depressive-like behavior observed with a minimal loss of locus coeruleus (LC) neurons following administration of 6-hydroxydopamine is associated with electrophysiological changes and reversed with precursors of norepinephrine. Neuropharmacology 101:76–86.

Tansey MG, Goldberg MS (2010) Neuroinflammation in Parkinson’s disease: its role in neuronal death and implications for therapeutic intervention. Neurobiol Dis 37:510–518.

Tong J, Hornykiewicz O, Kish SJ (2006) Inverse relationship between brain noradrenaline level and dopamine loss in Parkinson disease: a possible neuroprotective role for noradrenaline. Arch Neurol 63:1724–1728.

Valentino RJ, Van Bockstaele E (2008) Convergent regulation of locus coeruleus activity as an adaptive response to stress. Eur J Pharmacol 583:194–203.

Vazey EM, Aston-Jones G (2014) Designer receptor manipulations reveal a role of the locus coeruleus noradrenergic system in isoflurane general anesthesia. Proc Natl Acad Sci U S A 111:3859–3864.

Wakamatsu M, Ishii A, Ukai Y, Sakagami J, Iwata S, Ono M, Matsumoto K, Nakamura A, Tada N, Kobayashi K, Iwatsubo T, Yoshimoto M (2007) Accumulation of phosphorylated alpha-synuclein in dopaminergic neurons of transgenic mice that express human alpha-synuclein. J Neurosci Res 85:1819–1825.

Weinshenker D (2018) Long Road to Ruin: Noradrenergic Dysfunction in Neurodegenerative Disease. Trends Neurosci 41:211–223.

Yao N, Wu Y, Zhou Y, Ju L, Liu Y, Ju R, Duan D, Xu Q (2015) Lesion of the locus coeruleus aggravates dopaminergic neuron degeneration by modulating microglial function in mouse models of Parkinsons disease. Brain Res 1625:255–274.

Zarow C, Lyness SA, Mortimer JA, Chui HC (2003) Neuronal loss is greater in the locus coeruleus than nucleus basalis and substantia nigra in Alzheimer and Parkinson diseases. Arch Neurol 60:337–341.

